# Cortical excitation:inhibition imbalance causes abnormal brain network dynamics as observed in neurodevelopmental disorders

**DOI:** 10.1101/492108

**Authors:** Marija Markicevic, Ben D. Fulcher, Christopher Lewis, Fritjof Helmchen, Markus Rudin, Valerio Zerbi, Nicole Wenderoth

**Affiliations:** Neural Control of Movement Lab, HEST, ETH Zürich, Winterthurerstrasse 190, 8093 Zurich, Switzerland; School of Physics, The University of Sydney, NSW 2006, Australia; Brain Research Institute, University of Zurich, Winterthurerstrasse 190, 8057 Zurich, Switzerland; Neuroscience Center Zurich, University and ETH Zurich, Winterthurerstrasse 190, 8057 Zurich, Switzerland; Institute of Pharmacology and Toxicology, University of Zurich, Winterthurerstrasse 190, 8057 Zurich, Switzerland; Institute for Biomedical Engineering, University and ETH Zurich, Wolfgang-Pauli-Str. 27, 8093 Zurich, Switzerland

**Author notes:** V.Z and N.W share last authorship. **Corresponding author:** Dr. Valerio Zerbi, Neural Control of Movement Lab, Department of Health Sciences and Technology, ETH Zürich, Switzerland, Auguste-Piccard-Hof 1, 8093 Zurich.

**Keywords:** excitation:inhibition balance, fMRI, DREADD, hypoconnectivity, functional connectivity

## Abstract

Abnormal brain development manifests itself at different spatial scales. However, whether abnormalities at the cellular level can be diagnosed from network activity measured withfunctional magnetic resonance imaging (fMRI) is largely unknown, yet of high clinical relevance. Here we applied fMRI while using chemogenetics to increase the excitation-to-inhibition ratio (E:I) within cortical microcircuits of the mouse brain, thereby mimicking a putative mechanism of neurodevelopmental disorders including autism. Increased E:I caused a significant reduction of long-range connectivity, irrespective of whether excitatory neurons were facilitated or inhibitory Parvalbumin interneurons were suppressed. Training a classifier on fMRI signals, we were able to accurately classify cortical areas exhibiting increased E:I. This classifier was validated in an independent cohort of *Fmr1^y/-^* knock-out mice, a model for autism with well-documented loss of Parvalbumin neurons and chronic alterations of E:I. Our findings demonstrate a promising novel approach towards inferring microcircuit abnormalities from macroscopic fMRI measurements.

## 1. Introduction

Complex behavior results from the interplay of distributed yet anatomically connected neuronal populations which form brain-wide networks. Abnormal brain development as in Autism Spectrum Disorders (ASD) often affects the structural and functional connectivity within these networks at different spatial scales: at the cellular and (micro)circuit levels, neurodevelopmental disorders are associated with dysregulated synaptogenesis, cell excitability, synaptic transmission, and plasticity^1, 2^, as well as aberrant axonal migration and abnormal functioning of cortical (micro)circuits^3^; at the network level, long-range functional connectivity and macroscopic neuronal dynamics are altered^4, 5^. Mechanistically linking disease mechanisms across scales is an important goal for neuropsychiatry, particularly, to identify whether certain cellular disease pathways map onto specific alterations of large-scale network activity, which are accessible in human patients and might thus allow to infer the underlying pathobiology of an individual patient.

An abnormal increase of the excitation-to-inhibition (E:I) ratio within specific cortical circuits^6^ has been hypothesized as a putative disease mechanism underlying ASD pathology and its core behavioral symptoms^7, 8^. In transgenic mouse models of ASD, elevated E:I has been observed either because excitatory transmission is increased^9^, or because inhibitory transmission is reduced, for example, due to the loss of fast-spiking parvalbumin (PV) GABAergic interneurons^2, 10, 11^. However, it is unknown whether such an imbalance at the microcircuit level propagates across scales, and whether macroscopic markers of brain activity signal these abnormalities with anatomical specificity. Here we mimic neurodevelopmental pathology by acutely increasing microcircuit E:I within one brain region and test whether this well-controlled manipulation causes an anatomically specific change of neuronal dynamics and macroscopic connectivity of the mouse brain. We used chemogenetics^12, 13^ to manipulate cell activity of somatosensory cortex (SSctx) while simultaneously imaging brain activity using resting-state functional magnetic resonance imaging (rsfMRI). rsfMRI is a non-invasive method to measure spontaneous slow-frequency fluctuations of the blood oxygen level-dependent (BOLD) signal (an indirect estimate of neuronal activity) in the absence of a task and across the entire brain^14^. A growing body of evidence suggests that rsfMRI has great potential for translation, as it detects abnormalities in long-range network correlations in human patients as well as homologous networks in mouse models expressing similar transgenic mutations^15, 16, 17^. Here, we show that experimentally increasing E:I in primary somatosensory cortex (SSctx) by either (i) facilitating excitatory neurons, or (ii) suppressing inhibitory PV interneurons, both caused a *reduction* of long-range functional connectivity between SSctx and anatomically connected areas. Increasing E:I changed the dynamics of BOLD fluctuations, an effect that was most pronounced in the targeted SSctx. We used the BOLD timeseries measured during experimentally controlled chemogenetic suppression of PV interneurons to train a classifier to detect acutely over-excited brain areas. We applied this classifier to an independent cohort of *Fmr1^y/-^* knockout mice, a well-characterized model for ASD^4, 5^ which suffers from PV interneuron depletion and exhibits increased cortical E:I due to the loss of inhibitory inputs^18^. Importantly, the classifier identified specific brain regions in somatosensory and prefrontal cortex which exhibited an over-excitation phenotype based on their BOLD dynamics. This new finding indicates that mimicking microcircuit abnormalities in wildtype mice via chemogenetics allows us to build “computational sensors” for detecting E:I imbalances from non-invasive macroscopic rsfMRI measurements in a disease model. Our results demonstrate a promising novel approach towards inferring microcircuit abnormalities from macroscopic measurements which potentially opens new opportunities for investigating disease mechanisms across species.

## 2. Results

We mimicked ASD pathology by experimentally increasing the E:I balance in the right SSctx (Fig. S1). Adult mice were transfected with excitatory Designer Receptors Exclusively Activated by Designer Drugs (hM3Dq DREADDs), which were tuned to primarily target pyramidal neurons (wt-hM3Dq, *n* = 10). Control mice were sham operated (weight and age-matched littermate controls, *n* = 9; Table S1). Three to four weeks after surgery, mice were lightly anesthetized and electrophysiology or macroscopic brain activity was recorded before and after activating the DREADDs with a low dose of clozapine (0.01–0.03 mg/kg, i.v.; for a comparison to activating DREADDs with clozapine-N-oxide see Figure S2).

*In vivo* electrophysiology recordings in wt-hM3Dq mice (Fig. S1D) confirmed that activating the DREADDs increases net excitation within the targeted right SSctx but not in a control area, i.e., its homotopic counterpart (left SSctx). Multi-unit activity was recorded under isoflurane anaesthesia (0.5%) from multi-contact electrode arrays (Fig. 1A–D). During the baseline period (15 min) the average basal firing rate was stable in both brain areas. However, upon clozapine injection, the average firing rate in the right SSctx increased steadily over time while activity in the left SSctx remained virtually unchanged. 30 min after injection, the difference in averaged multi-unit activity between left and right SSctx reached 11% (Fig. 1A–D).

**Figure 1.**
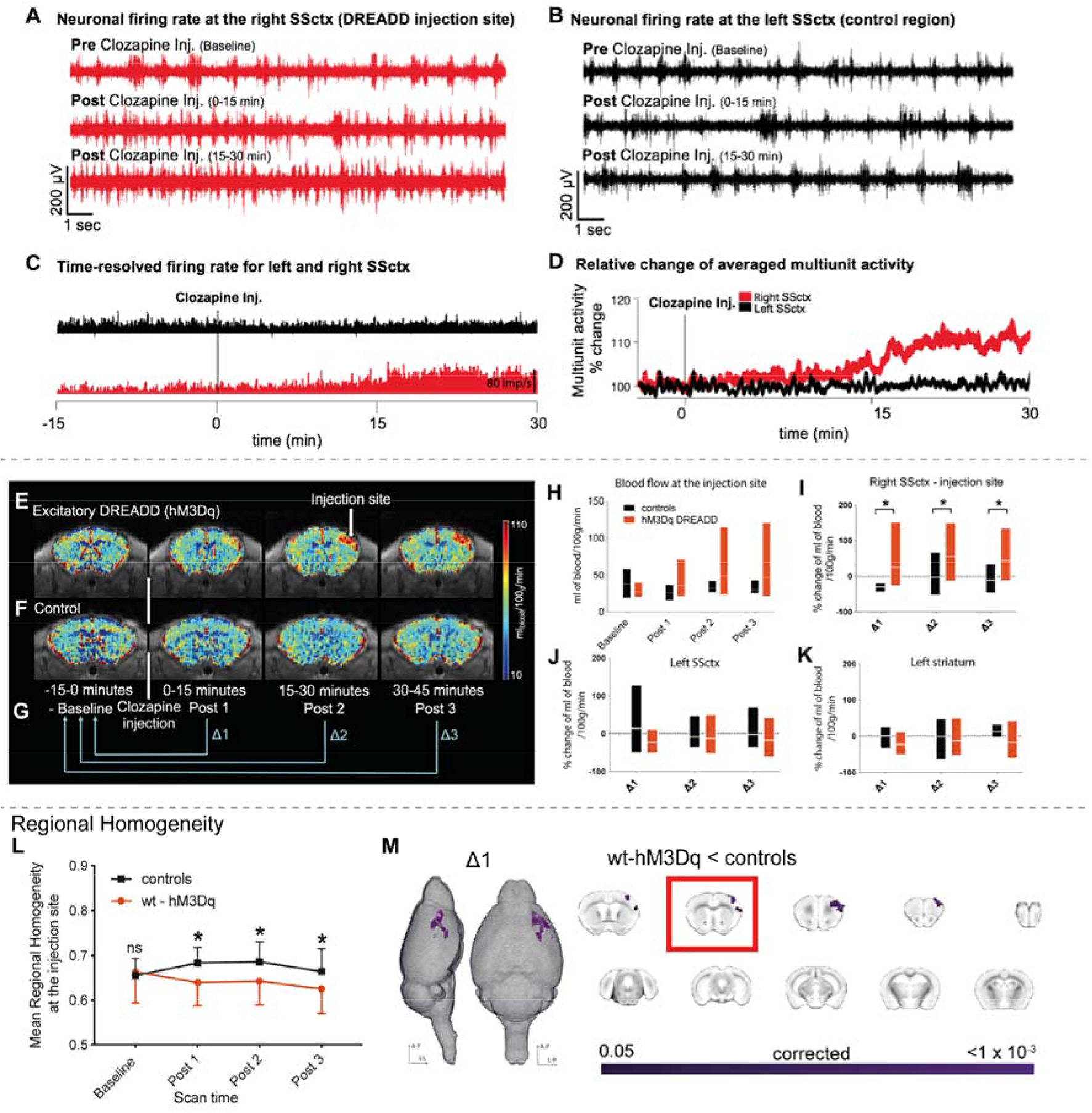
Local changes in neural activity induced by activating hM3Dq. **A)** Neuronal firing rate before and after clozapine injection in the right SSctx, indicating a steady increase in firing rate upon DREADD activation. **B)** As A but for the left SSctx, (control area) indicating no change in the neuronal firing rate after clozapine injection. **C)** Time-resolved firing rate in millisecond bins for the right (red) and left (black) SSctx. A steady increase in firing rate occurs in the right SSctx once clozapine is injected, while no change occurs in the left SSctx. **D)** Relative change of averaged multiunit activity recorded in the right and left SSctx. **E), F)** Cerebral blood flow maps were collected every 15 min wt-hM3Dq and control mice, respectively. **G)** Clozapine is injected 15 min after the start of experiment. These 15 minutes are referred as the baseline period (–15-0 min). The rest of the experiment is 45 minutes long and divided into 3 periods i.e., Post 1 (0-15 min), Post 2 (15-30 min), and Post 3 (30-45 min). For all analyses performed in this experiment (unless otherwise stated), Post data was expressed relative to baseline (Post 1/Post 2/Post 3 – baseline) and is referred to as Δ1, Δ2, Δ3, respectively. **H)** Comparison of blood flow (ml of blood/100g/min) between controls and wt-hM3Dq mice over time (measured at the injection site). **I)** Percentage change in blood flow over time between the controls (n=7) and wt-hM3Dq (n=7) mice (measured at the injection site). Linear mixed models indicate a significant main group effect indicating an increase in blood flow in the wt-hM3Dq mice F_2,21_ = 5.657; p=0.04). **J)** Percentage change in blood flow over time measured in left SSctx. **K)** Percentage change in blood flow over time measured in left lateral striatum. **L)** Average regional homogeneity (ReHo) change over time at the injection site. Repeated measures ANOVA (control (n=13); wt-hM3Dq (n=14)) showed a significant interaction between scan time and groups F_1.93,75_ = 11.8; p<0.001). **M)** Significant decrease in ReHo for Δ1 depicted as a 3D image and in coronal slices (TFCE-corrected). The slice marked with a red rectangle shows the injection site.

Next, we used non-invasive fMRI to test whether activation of hM3Dq-DREADD would elicit anatomically specific changes in cerebral blood flow (CBF) which reflects glucose metabolism and has been shown to be an indirect marker of neural activity at the cell population level^19, 20^. After administering clozapine, wt-hM3Dq mice showed an increase in CBF around the injection site (Fig. 1E,H). The increase was significantly higher than in sham-operated mice (controls) scanned with identical procedures for all time points up to 45 min (Linear mixed model: F_2,21_ = 5.657, *p* = 0.04, Fig. 1F,I). No changes in CBF were observed in other brain regions, i.e., the contralateral somatosensory cortex and ipsilateral striatum (Fig. 1J, K). Together our electrophysiological and CBF results show that activating the DREADDs caused microcircuit excitation which was anatomically specific to the injected area.

### Activating hM3Dq-DREADD decreases short-range connectivity selectively within somatosensory and somatomotor cortex

We next tested whether hM3Dq-DREADD also caused a local disruption of functional connectivity (FC) measured with rsfMRI. To this end, we calculated the regional homogeneity (ReHo) index, a commonly used metric to assess the synchronization of BOLD fluctuations across neighboring voxels. We chose ReHo because it has been shown to reflect alterations occurring in various psychiatric diseases including depression, schizophrenia, and autism^21, 22^. Activation of hM3Dq-DREADD significantly reduced regional homogeneity only at the injection site when compared to controls (blue cluster in Fig. 1L, M; *p* < 0.05, TFCE-corrected). These changes occurred within the first 15 minutes after clozapine injection and were maintained for the whole scan period (Fig. S3A). No other area exhibited similar changes.

### HM3Dq DREADD reduces functional connectivity between somatosensory cortex and monosynaptically connected cortical areas

Does an increase in local microcircuit excitation also affect long-range FC as quantified by correlating spontaneous BOLD fluctuations between areas? To address this question, we measured FC between 130 brain regions as identified by the Allen Brain Reference Atlas Common Coordinate Framework (CCFv3) at every one-minute intervals (see Methods). We then extracted the effect size using the standardized difference between the group means (Cohen’s *d,* wt-hM3Dq vs control mice) between each pair of regions. During the first 15 minutes (baseline), Cohen’s *d* varied on average between −0.2 and +0.2 (null-to-small effect), and did not exhibit an appreciable spatial or temporal pattern. However, immediately after clozapine injection, we noticed a rapid reduction in FC between a number of connections in the somatosensory and somatomotor cortices, which remained hypoconnected for the whole scan session (Fig. 2A,B).

**Figure 2.**
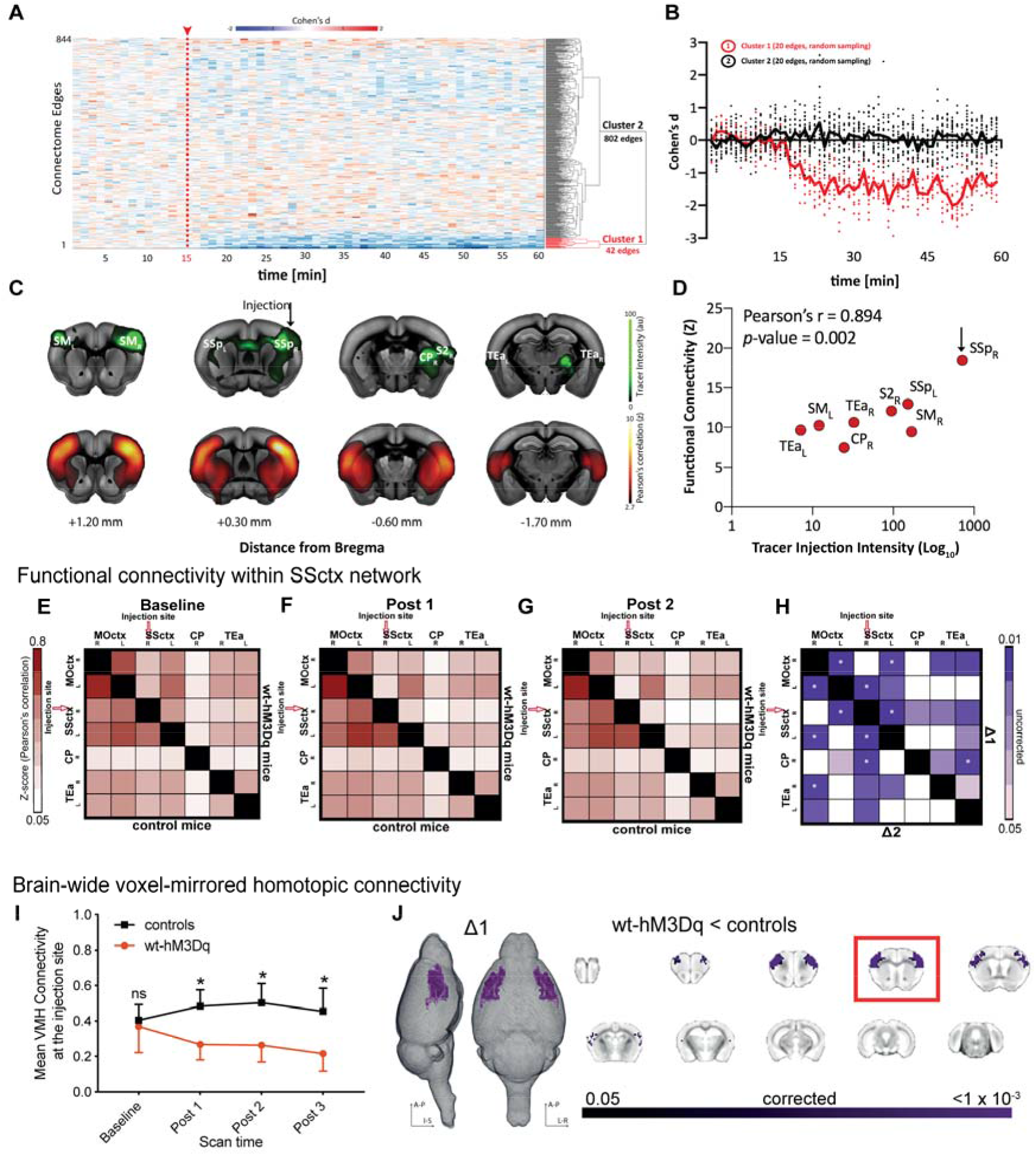
Long-range connectivity changes induced by activating hM3Dq. **A)** Effect-size analysis (Cohen’s d) of functional connectivity is calculated for single-edges (n=844). Only cluster 1 shows a decrease in the effect size after Clozapine injection. **B)** Averaged effect-size (Cohen’s d) over time for 20 edges of Cluster 1 and 20 randomly selected edges of Cluster 2. **C)** Coronal brain slices depicting tracer data after unilateral injection in SSctx (green, obtained from Allen Mouse Brain Connectivity Atlas, Injection ID: 114290938) and averaged voxel-wise Pearson’s correlation (Fisher’s z-transform) during the baseline period from a seed in the right SSctx. **D)** Pearson’s correlation between functional connectivity of the target region (right SSctx) and tracer injection intensity obtained from images depicted in C. **E) F) G)** Matrices represent averaged and z-transformed Pearson correlation of all control animals (lower triangular matrix) and all wt-hM3Dq animals (upper triangular matrix) for the baseline period, Post 1 and Post 2, respectively. **H)** The upper triangle of the matrix shows a reduction in FC in wt-hM3Dq (n=14) mice compared to controls (n=13) during Δ1 (i.e. Post1-baseline), while the lower triangle shows the same set of comparisons in Δ2 (i.e., Post2-baseline). Color coding reflects randomized permutation testing, p < 0.05, uncorrected. Asterisks indicate significant differences that survived FDR correction for multiple comparisons. MOctx – Somatomotor cortex, SSctx – somatosensory cortex, CP –caudoputamen and TEa – temporal association cortex, R – right, L – left. **I)** Change over time of the mean brain-wide VMHC (symmetric connectivity) at the injection site. Repeated measures ANOVA (controls (n=13); wt-hM3Dq (n=14)) showed a significant interaction between the scan time and groups (F_1.67,41.9_ = 21.18; p < 0.001). **J)** Significant decrease (TFCE-corrected) in symmetric connectivity depicted on 3D images and coronal slices for the Δ1 time point. The slice marked with a red rectangle is the injection slice.

We next quantified specific changes in FC within the somatosensory network, i.e. areas which are monosynaptically connected to the chemogenetic target, right SSctx according to the Allen Mouse Brain Connectivity Atlas^23^. This anatomically defined network includes contralateral primary SSctx, bilateral somatomotor cortex (MOctx), bilateral association areas (TEa) and ipsilateral caudatum putamen (CP) (Fig. 2C). We found high FC between all of these structures at baseline (Fig. 2C), and a strong structure–function correspondence (Fig. 2D). To increase the power of our analysis, the BOLD time series of the somatosensory network regions were binned into four intervals of 15 minutes each. Pearson’s correlations were calculated between regions and are summarized in FC matrices (Figs 2E, F, G, and Fig. S4A). At baseline, FC did not show differences between groups (Fig. 2E). By contrast, after clozapine injection (Fig. 2F–H) we observed a reduction in FC between right SSctx and nearly all other areas of the network in wt-hM3Dq mice, including cortico-striatal circuitry. Significant group differences (permutation testing, *p* < 0.05, FDR corrected) were found between right SSctx and left SSctx/left MOctx, as well as between left and right MOctx, and were maintained for the duration of the scan session (Fig. 2H, Fig. S4B).

We further complemented our analysis by assessing the similarity between the BOLD time series of symmetric voxels in the left and right hemisphere. This index has been shown to identify pathological connectivity patterns with high sensitivity as compared to other FC metrics^24^. In line with our previous findings, injecting clozapine caused a significant reduction of symmetric connectivity in wt-hM3Dq mice but not in controls which occurred only between the left and right primary SSctx, and left and right MOctx (*p* < 0.05, TFCE corrected) (Fig. 2I, J). In summary, our analyses demonstrate that increasing local cortical excitation causes a reduction in local connectivity that propagates into long-range FC reduction exclusively within the targeted network.

The above results were obtained using human Synapsin-1 as the promoter for the viral injection. However, it has been argued that, depending on the titer, transfection might not be specific for pyramidal neurons. To address this concern, we repeated the above rsfMRI experiment in a separate cohort of wildtype mice (n=13) using the CAMKII promoter for AAV hM3Dq-DREADD delivery (Fig. S5A-C). Importantly, we replicated our main results which consistently confirms that increasing E:I by enhancing excitatory synaptic transmission causes a robust reduction of FC within an anatomically connected network.

### Changing E:I balance by inhibiting GABAergic parvalbumin neurons

Changes in E:I have been proposed as a principle mechanism that underlies symptoms of neurodevelopmental disorders, irrespective of whether primarily excitatory or inhibitory synapses are affected. Next, we tested whether suppressing the activity of GABAergic PV interneurons — as reported in several mouse models for ASD^2, 11, 25^ — would cause similar macroscopic hypo-connectivity as observed above. Nineteen Pvalb^tm1(cre)Arbr^ (PVCre) were transfected with an inhibitory hM4Di-DREADD. Clozapine activation reduced PV cell activity which resulted in over-excitation of right SSctx but not a control region. Accordingly, average firing rate in right SSctx went up by 20% following clozapine injection (Fig. 3A,B and Fig S6A,B) and local cerebral blood flow increased moderately (repeated measures ANOVA: F_1.4,17.9_=2.45; *p*=0.1; Fig. 3C). Our rsfMRI analysis confirmed the expected reduction in local synchronization around the injection site (Fig. 3D) and also marked under-connectivity which was most pronounced between right and left SSctx (Fig. 3E-H) in PVCre-hM4Di mice as compared to controls. In summary, our results reveal that shifting E:I toward excitation by inhibiting PV interneurons leads to a reduction of local and long-range FC, which is qualitatively similar to that observed during direct system excitation.

**Figure 3.**
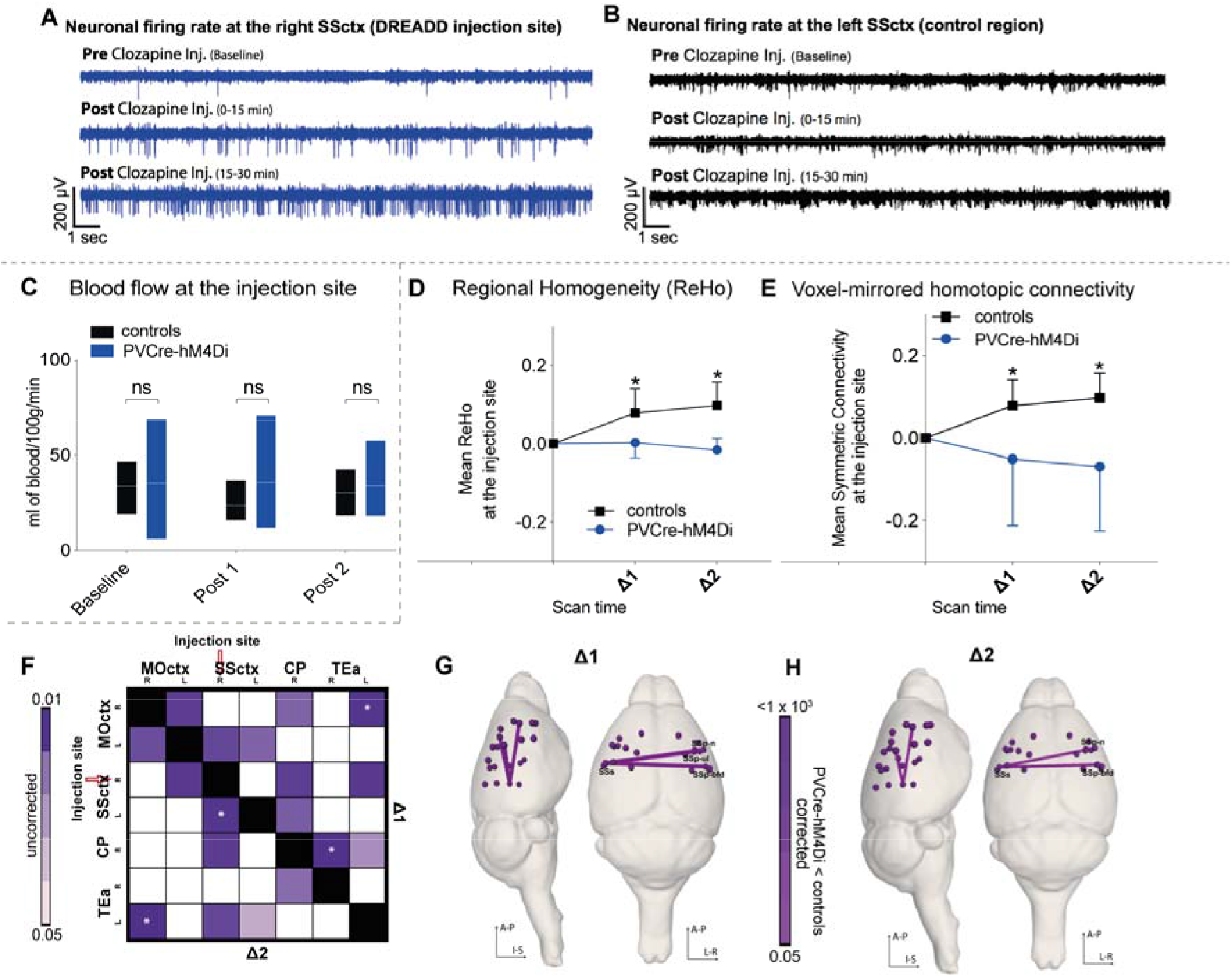
Changes induced by activating hM4Di in PVCre mice. **A)** Neuronal firing rate at the right SSctx DREADD injection site of the PVCre-hM4Di mice before and after clozapine injection, indicating a steady increase in firing rate on DREADD activation. **B)** Similar to figure A but for right caudate putamen, indicating no change in the neuronal firing rate after clozapine injection. **C)** Comparison of blood flow (ml of blood/100g/min) between controls (n=7) and PVCre-hM4Di (n=13) mice over time (measured at the injection site). No significant differences observed between groups (repeated measures ANOVA: F_1.4,17.9_=2.45; p=0.1). **D)** Normalized average regional homogeneity (ReHo) change over time at the injection site indicating significant differences between groups (controls: n=13; PVCre: n=19; repeated measures ANOVA: F_1,25_=14.01; p=0.001). Post hoc paired t-test revealed a significant group difference at Δ1 ((t_12_=-3.60; p=0.004) and at Δ2 (t_12_=-8.05; p < 10^^4^)). **E)** Normalized averaged symmetric connectivity over time at the injection site. Repeated measures ANOVA (controls (n=13); PVCre-hM4Di (n=19)) showed a significant interaction between scan time and group F_1.2, 21.9_=8·21; p = 0.007). Post hoc paired t-test revealed a significant group difference at Δ1 (p=0.047) and at Δ2 (p = 0.004). **F)** Seed to seed analysis indicates reduced FC (randomized permutation testing, p < 0.05, uncorrected) between controls (n=13) and PVCre-hM4Di (n=19) mice. **G) H)** Whole-brain connectome analysis shows a significant interhemispheric reduction between somatosensory cortices for Δ1 and Δ2 between PVCre-hM4Di (n=14) mice and controls (n=13). Regions affected are as follows: SSp-m: Primary Somatosensory Area, mouth; SSp-ul: Primary Somatosensory Area, upper limb; SSs: Supplementary somatosensory area; SSp-bdf: Primary Somatosensory Area, barel field; SSp-n: Primary Somatosensory Area, nose.

### Altered E:I balance changes local BOLD dynamics

Our previous analyses demonstrated that increasing E:I in right SSctx affects its FC with other brain regions. We next aimed to understand how microcircuit E:I shapes the fluctuations of spontaneous BOLD regional activity. As depicted in Fig. 4A, we focused our analysis on three regions of interest: (i) right SSctx (the injection site); (ii) left SSctx (its homotopic counterpart); and (iii) left visual cortex (a control region). The dynamics of BOLD time series were characterized by an interdisciplinary library of over 7000 features (using the software package *hctsa*^26^), which capture distinct properties of the BOLD signal, including spectral power in different frequency bands, temporal entropy, and many more^26, 27, 28^ (Fig. 4B, see Methods for details).

**Figure 4:**
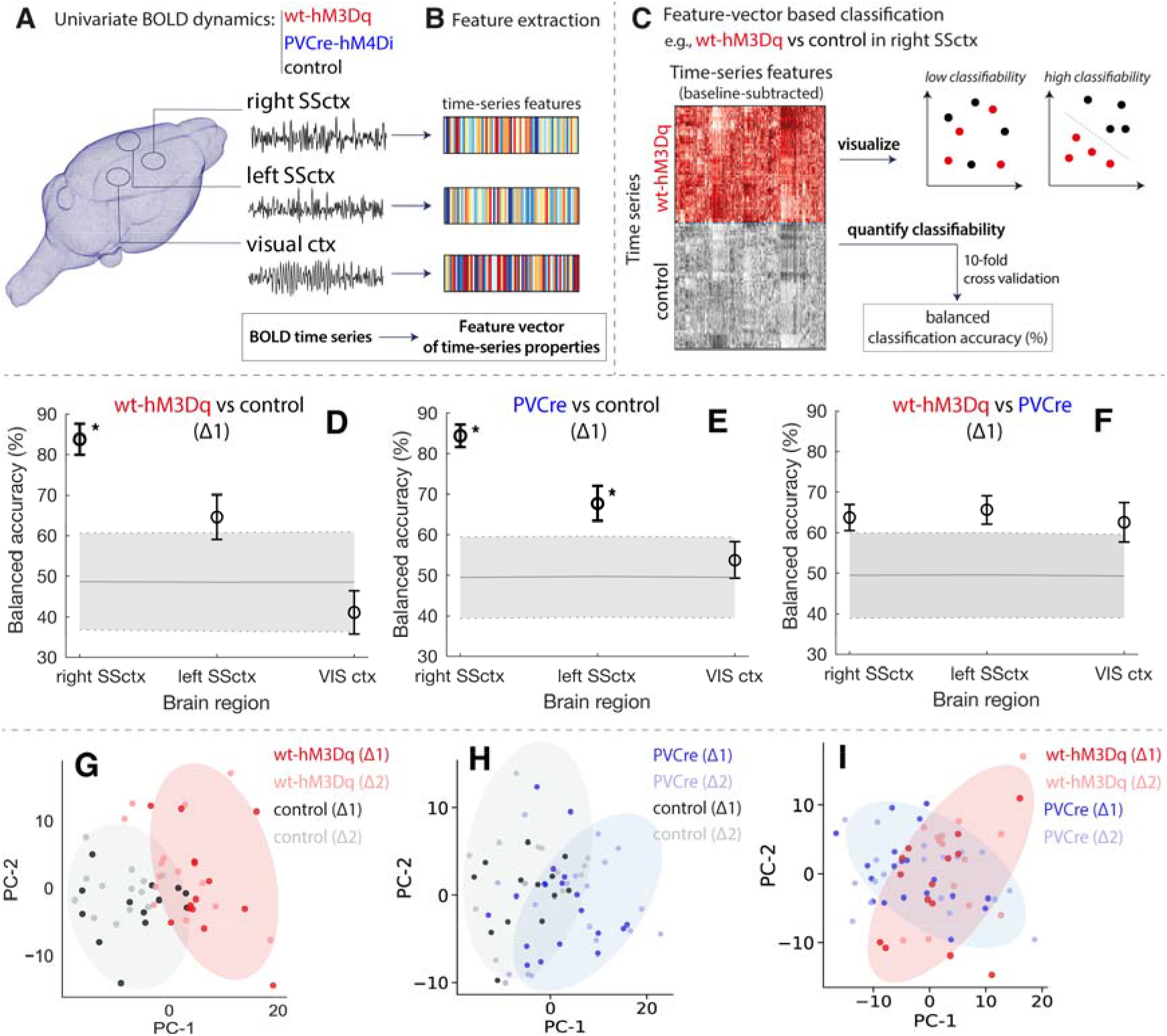
DREADD manipulations lead to characteristic changes in univariate BOLD dynamics. Classification of BOLD signal dynamics in three brain regions (the injected right SSctx, left SSctx, and VISctx) across three conditions (wt-hM3Dq, PVCre-hM4Di, and control). A schematic of the approach is depicted in **A–C: A)** BOLD dynamics are measured from each brain region as a univariate time series (a 15 min time series per time window and experiment), which was **B)** converted to a set of properties (a ‘feature vector’) using hctsa. **C)** For a given region and pair of classes, we used the features of each time series (relative to baseline) as the basis for classification, which was visualized using a low-dimensional principal components projection and quantified as the 10-fold cross-validated balanced classification accuracy (%). Classification results in each brain region at Δ1 are shown for: **D)** wt-hM3Dq (n=14) versus control (n=13), **E)** PVCre-hM4Di (n=10) versus control (n=13), and **F)** wt-hM3Dq (n=14) versus PVCre-hM4Di (n=10) revealing significant discriminability in the right SSctx for wt-hM3Dq and PVCre-hM4Di versus control (permutation test, p < 0.05, annotated as ‘* ‘). There is a consistent trend of high discriminability in the injected region (right SSctx), followed by the contralateral region (left SSctx), and lowest discriminability in the control region (VISctx). BOLD time series in right SSctx are visualized in two-dimensional principal components spaces in **G)–I)** for the same three pairs of classes as in **D)–F)**. Time series with similar properties are close in the space, revealing a visual depiction of the discriminability of wt-hM3Dq and PVCre-hM4Di relative to control (G, H), but a relative lack of discriminability for wt-hM3Dq versus PVCre-hM4Di (I). Shaded ellipses (Δ1) have been added to guide the eye, and time series from each class are labeled for Δ1 and Δ2.

Based on the change of these features from baseline to activated DREADDs, we trained linear Support Vector Machine (SVM) classifiers to distinguish: (i) wt-hM3Dq versus controls, (ii) PVCre-hM4Di versus controls, and (iii) wt-hM3Dq versus PVCre-hM4Di, as shown in Fig. 4C.

Our classification results reveal that BOLD dynamics of the injected right SSctx at Δ1 are differentiated from control mice in both wt-hM3Dq (Fig. 4D) and PVCre-hM4Di (Fig. 4E) with high accuracy (≥84% balanced accuracy after 10-fold cross-validation), significantly exceeding chance levels (p ≤ 1 × 10^-3^, permutation test). Qualitatively similar results were obtained at Δ2 (see Supplementary Results). DREADD activation had a smaller effect on the accuracy of classifying BOLD dynamics in the left SSctx for both wt-hM3Dq, 64% *(p* = 0.1), and PVCre-hM4Di, 69% *(p* = 0.03), and was weakest for the control region, VISctx, in both wt-hM3Dq, 43% *(p* = 0.7), and PVCre-hM4Di, 54% *(p* = 0.3).

To aid visualization, we constructed principal component projections of each dataset (including data at Δ2), shown for wt-hM3Dq versus control and PVCre-hM4Di versus control in Figs 4G and 4H, respectively. These plots reveal a clear distinction between DREADD-activated and control mice on the basis of their BOLD dynamics in right SSctx.

Since wt-hM3Dq and PVCre-hM4Di increase E:I through different mechanisms, we next investigated whether these two manipulations could be distinguished from each other on the basis of their BOLD dynamics during Δ1. As shown in Fig. 5F, BOLD dynamics between wt-hM3Dq and PVCre-hM4Di are only weakly distinguishable in the injected region, 64% *(p* = 0.09), contralateral region, 66% *(p* = 0.06), and control region, 63% *(p* = 0.1), and the clear discrimination of BOLD dynamics in principal components space (Fig. 4G,H) is starkly absent (Fig. 4I, see strongly overlapping distributions). To further compare the effect of wt-hM3Dq and PVCre-hM4Di manipulations on BOLD dynamics, we computed a similarity measure between changes in BOLD time-series properties caused by both manipulations relative to controls (as a Spearman’s *ρ* in right SSctx and left SSctx relative to the control region across all time-series features, see Fig. S9). We found overall agreement between wt-hM3Dq and PVCre-hM4Di in right SSctx (*ρ* = 0.59) and left SSctx (*ρ*= 0.47), suggesting that the temporal properties of BOLD dynamics are more sensitive to the net over-excitation common to both DREADD cohorts, than to the underlying synaptic transmission pathway within the cortical microcircuit (see Supplementary Material for an analysis of BOLD time-series features that are individually informative of E:I).

**Figure 5:**
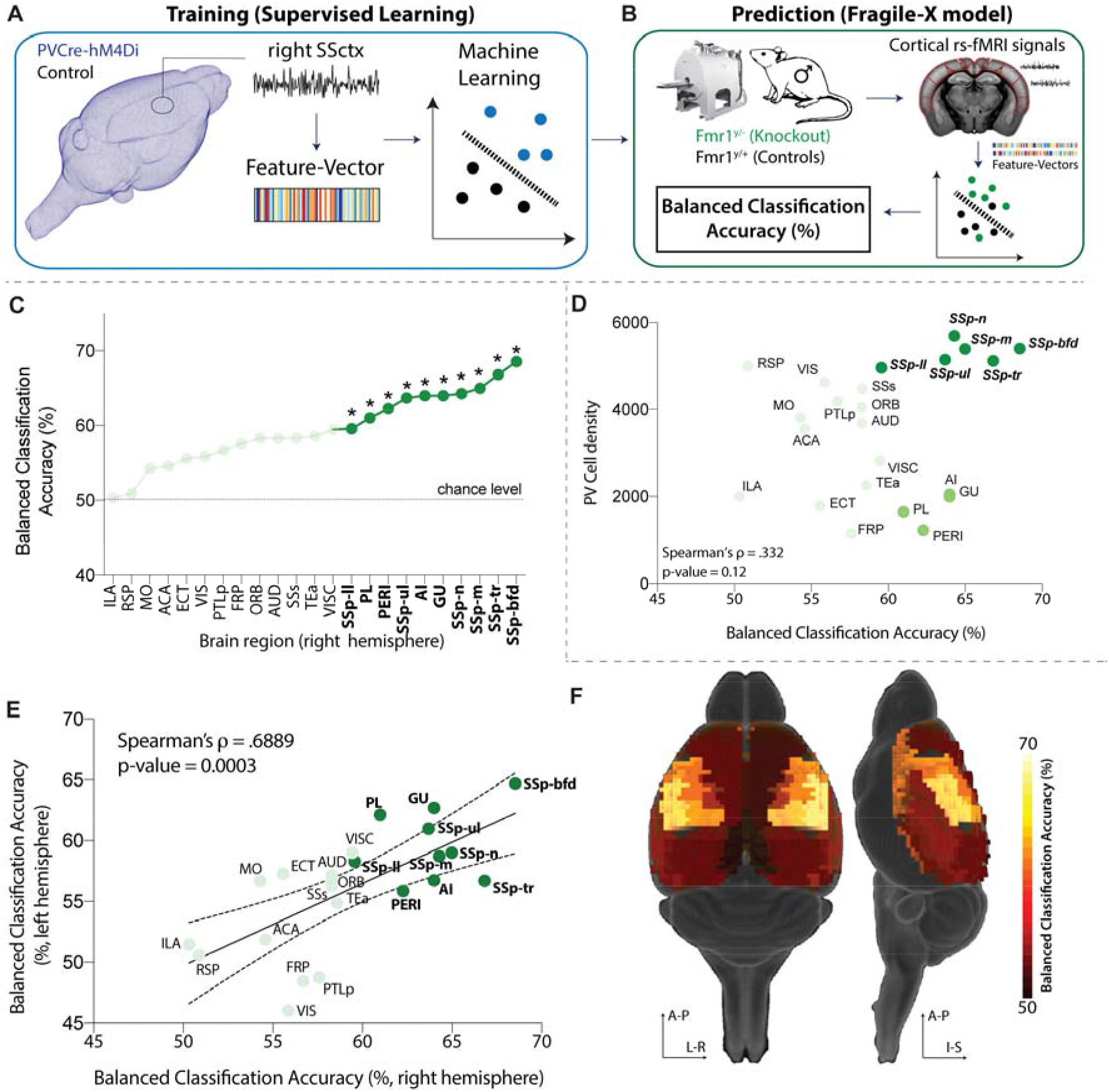
Fmr1^y/-^ exhibit similar local BOLD dynamics as DREADD-manipulated mice. **A)** The classifier was trained on features of right SSctx BOLD time series to distinguish PVCre-hM4Di from control mice. **B)** The trained classifier was used to predict the identity of cortical BOLD time series measured from Fmr1^y/-^ (knockout) and Fmr1^y/^+ (control) mice, identifying predictions of PVCre-hM4Di as Fmr1^y/-^. Predictability in each brain region was assessed as balanced classification accuracy (%); p-values were estimated using a permutation test. **C)** Classification results are shown for the right hemisphere, highlighting regions with significant classification accuracy (p<0.05, FDR-corrected) as bold, dark green, and marked with asterisks. **D)** Scatter plot of balanced classification accuracy and PV cell density. **E)** Balanced classification accuracy is plotted for each region in the right and left hemispheres, exhibiting a strong positive correlation, Spearman’s ρ = 0.69 (p = 3 × 10^4^). **F)** Balanced classification accuracy visualized on a 3D brain.

### *Fmr1^y/-^* exhibit similar local BOLD dynamics to DREADD-manipulated mice

Our previous analysis yielded a classifier for predicting an acute microcircuit E:I increase from macroscopic BOLD dynamics. Next, we asked whether such a classifier can also detect abnormal BOLD dynamics in a transgenic disease model of ASD. We used rsfMRI data from *Fmr1^y/-^* knock-out mice that mimic Fragile X syndrome, a neurodevelopmental disorder characterized by cognitive impairment, epileptic seizures, physical abnormalities and hypersensitivity. Previous studies in *Fmr1^y/-^* mice report a reduction in cortical PV cell density, specifically within somatosensory region, followed by cortical hyperexcitability^9, 11, 29, 30^ These findings indicate that the disease mechanism of *Fmr1^y/-^* mice affects the net excitability of cortical microcircuits in a similar manner as chemogenetically suppressing PVCre activity, but it is unknown whether these circuit-level alterations cause analogous alteration of macroscopic rsfMRI dynamics. To test this hypothesis, we trained a classifier for PVCre-hM4Di rsfMRI time series in the right SSctx during Δ1 as above (Fig. 4E; 5A). We then applied this classifier to rsfMRI data from 46 cortical ROIs (23 from each hemisphere) in *Fmr1^y/-^* mice (n = 44) and wildtype littermates *(Fmr1^y/^+, n* = 39) and assigned predicted labels to each brain region of *Fmr1^y/-^* and *Fmr1^y/^+* independently (Fig. 5B). As shown in Fig. 5C, regional BOLD dynamics of *Fmr1^y/-^* versus wildtype controls could be significantly classified in 10 different areas of the right hemisphere (balanced classification accuracy, *p* < 0.05, FDR-corrected) using a classifier trained on PVCre-hM4Di data. Corresponding to the brain area in which the DREADDs classifier was trained, regions of the SSctx displayed the highest classifiability. Similar results were obtained in the left hemisphere, resulting in a Spearman’s *ρ* = 0.69 *(p* =3 x10^-4^) between the left and right hemispheres (Fig. 5E, F). We next tested whether regional variation in classification accuracy was related to differences in PV cell density using data from *qBrain*^31^. As shown in Fig. 5D, classification accuracy is only weakly correlated with PV cell density, *ρ* = 0.33 *(p* = 0.1), but is highest in SSp regions with the highest PV cell density (dark green in Fig. 5D). Interestingly, the classifier identified several prefrontal areas (bright green in Fig. 5D) which exhibited abnormal rsfMRI dynamics according to the classifier despite low PV cell density. This finding is in line with former reports of abnormal prefrontal cortex function in *Fmr1^y/-^* mice^32, 33^. Our results demonstrate that a classifier trained on a controlled chemogenetic manipulation causing an acute shift of the E:I balance is generalizable to *Fmr1*^y/-^, indicating a strong degree of similarity between cortical BOLD dynamics resulting from PVCre-hM4Di and *Fmr1^y/-^* knockout.

## 3. Discussion

It has been a longstanding question how disease mechanisms at the cellular level map onto abnormal dynamics at the level of macroscopic brain networks. Here we show in four independent cohorts that mimicking neurodevelopmental pathology by acutely increasing the microcircuit E:I balance in somatosensory cortex via chemogenetics causes distinct changes of macroscopic brain dynamics and severe functional under-connectivity between the manipulated region of interest (SSctx) and monosynaptically connected brain areas. These macroscopic changes were observed irrespective of whether we enhanced excitatory transmission or selectively suppressed inhibitory PV neurons of the same cortical circuits. Our results constitute the first direct evidence that macroscopic BOLD dynamics reflect the state of underlying cortical microcircuits. Furthermore, we show that different cellular mechanisms can cause common pathophysiological abnormalities and converge on highly similar alterations of neural dynamics at the network level. In particular, upon activating the DREADDs there was a marked reduction of FC which was temporally locked to our stimulation paradigm and spatially specific to the injected area and its monosynaptically connected neighbors. Reduced FC has been frequently reported in human fMRI studies of neurodevelopmental disorders, such as autism and schizophrenia^34, 35, 36, 37, 38^, as well as in relevant mouse models of these disorders (*CNTNAP2, Fmr1^y/-^, En2*)^4, 39^. Many of these mouse models have been shown to exhibit an increased E:I^3, 6, 9^. Additionally, they suffer from reduced PV interneuron expression^2, 10^. Depletion of this specific cell type has been hypothesized to cause aberrant patterns of neural activity and connectivity^2, 11^, as fast-spiking PV interneurons target the proximal regions of pyramidal cells^40, 41^ and synchronize firing within pyramidal cell populations^42, 43, 44, 45, 46, 47^. This hypothesis has been recently confirmed by Marissal and colleagues, who found that DREADD-induced suppression of PV activity causes the disinhibition of pyramidal neurons and leads to mass desynchronization of spiking activity^48^. Congruent with these previous empirical findings and recent theoretical models^49^, our data suggest that inhibiting PV neurons reduces the *synchronization* of neuronal ensembles as indicated by the reduction of FC, despite higher net cortical activity.

Interactions among different types of neurons within a microcircuit should be balanced for proper functioning. Lee and colleagues^49^ have recently built a computational model of the sensory cortex microcircuit and demonstrated the importance of connectivity homeostasis between neuronal types (i.e. inhibitory and excitatory cells) for producing synchronized oscillatory behavior with maximum frequency tuning capacity. Any deviation from this optimal configuration, such as over-excitation of pyramidal neurons, breaks this synchrony and impairs the communication among neuronal ensambles, which at the macro-scale may be perceived as a reduction in FC. In line with this theoretical framework, we observed a reduction in FC when DREADDs were used to activate pyramidal neurons. We replicated this principle finding across three independent cohorts, using either clozapine or CNO to activate the DREADDs (see supplementary discussion) and using either human Synapsin (hSyn) or CAMKII as a promoter in the AAV constructs. Both promoters generate a clear bias towards gene expression in excitatory cortical neurons (up to 85%) even though they might also transfect a small number of inhibitory neurons. Importantly, our electrophysiological results confirmed that activating excitatory DREADDs in the wt-hM3Dq mice caused a significant net increase of neuronal firing rate and our CBF measurements confirmed that such an increase was anatomically specific and most pronounced with the targeted SSctx. Activating the excitatory DREADDs caused overall a stronger reduction of FC in wt-hM3Dq mice than in PVCre-hM4Di mice, which might result from a higher number of transfected neurons: mouse cortex contains 12% GABAergic neurons, of which approximately one quarter are PV^50^ resulting in a rather low absolute number of transfected neurons and smaller DREADD induced effects^51^.

Our results also characterize a potential role of local E:I balance in shaping the BOLD dynamics of individual brain regions. Specifically, we showed that the chemogenetic manipulation of the cellular excitability causes specific changes of local BOLD features that we could distinguish from neurotypical BOLD dynamics with high accuracy (balanced accuracy >80%) and anatomical specificity, irrespective of whether we increased excitatory transmission (wt-hM3Dq cohort) or suppressed inhibition (PVCre-hM4Di cohort). Importantly, the E:I BOLD signature learned from acutely modulating cell activity, could be detected in two independent cohorts of *Fmr1^y/-^* mice, a well-characterized mouse model of ASD. We chose *Fmr1^y/-^* mice for external validation because they are known to have deficits in local feedback inhibition due to reduced PV interneuron density^29, 52^. This leads to increased excitability and decreased synchrony of pyramidal cell firing through gamma oscillations^9, 11^ as well as substantially reduced FC in corticocortical and corticostriatal circuits^4, 53^. Interestingly, above chance-level detection accuracies of BOLD dynamics characterizing overexcitation in *Fmr1^y/-^* mice were found in regions which normally exhibit high PV cell densities^29^ providing further indirect evidence that the specific features of abnormal BOLD dynamics are related to the depletion of PV neurons in this mouse model.

Our findings demonstrate that a chemogenetically inspired and experimentally validated classifier can be used as a “computational sensor” which uses BOLD dynamics to infer whether a specific brain area exhibits increased microcircuit E:I for example, due to abnormal PV neuron activity.

It is unclear whether this discovery is generalizable to the entire cortex since changes in microstructural properties such as cytoarchitecture, E: I, size, density and laminar distribution of distinct cell types vary continuously through mouse cortex^54, 55^. These variations in spatial gradients shape functional specialization along the cortex and influence the intrinsic time scale that can be measured in BOLD dynamics in different cortical areas^28^. Nevertheless, we show above chance-level detection accuracy in the prefrontal cortex of *Fmr1^y/-^* mice that are also attributed to an increased E:I balance from previous studies^32^. These results provide a new framework for mechanistically linking microscale circuit dynamics to macroscopic fMRI-based brain activity at rest. Thus far, this approach is unique because it uses causal microcircuit alterations to train a classifier that can detect similar alterations from distinct datasets obtained from large-scale imaging techniques. Our framework goes beyond describing long-range connectional aberrations caused by neurodevelopmental disorders and aims at providing deeper insights into how distinct microcircuit dynamics relate to large-scale, non-invasive imaging measurements which can be obtained across different species, including in human patients. Better linking macroscopic alteration to the underlying neurobiology is a pre-requisite for advancing neuroimaging as a tool in the context of clinical decision making.

## 4. Materials and Methods

All experiments and procedures were conducted following the Swiss federal Ordinance for animal experimentation and approved by Zurich cantonal Veterinary Office (ZH238/15). C57BL/6 mice were obtained from Charles River Laboratories (Germany), while Pvalb^tm1(cre)Arbr^ (PVCre) mice were derived from in-house breeding (first generation obtained from the Jackson Laboratory). All mice were kept in standard housing under 12h light/dark cycle with food and water provided *ad libitum* throughout the whole experiment. A total of 29 male C57BL/6 mice were used in the experiment, aged 9 ± 1 weeks and weighing 25.1 ± 1.8gr (mean ± SD) at the day of the surgery. A total of 19 PVCre mice were utilized in this experiment, aged 11 ± 1.6 weeks and weighing 26.1± 3.8gr at the day of the surgery.

### Procedures for wildtype (hM3Dq) mice study

#### Transfection procedures for wildtype (wt-hM3Dq) and CAMK-hM3Dq mice

Each mouse was initially anesthetized using a mixture of midazolam (5mg/ml; Sintetica, Switzerland), fentanyl (50mcg/ml; Actavis AG, Switzerland) and medetomidine (1mg/ml; Orion Pharma, Finland). Upon anesthesia induction, mice were placed on a heating pad and the temperature was kept at 35°C (Harvard Apparatus, USA). Following shaving and cleaning, an incision along the midline of scalp was made. The right primary somatosensory cortex was targeted at the coordinates of +0.5 mm AP (anterior-posterior), −3.0 mm ML (medio-lateral) and −2.25 mm DV (dorso-ventral) relative to Bregma using a drill and microinjection robot (Neurostar, Germany) with a 10 ul NanoFil syringe and 34Ga bevelled needle (World Precision Instruments, Germany). 950 nl of ssAAV8-hSyn-hM3Dq-mCherry (n=13) of a physical titer ≥ 5.4 x 10^12^ vg/ml (*vector genomes; fluorometric quantification) was injected at the rate of 0.06 ul/min. The virus was provided by Viral Vector Core Facility of the Neuroscience Centre Zurich (http://www.vvf.uzh.ch/en.html). Upon the injection, the needle was left in place for 10 min and then slowly withdrawn. C57BL/6 control mice (n=9) underwent the same surgical procedures, where the needle was kept in place for 5min but without any viral injections. Subsequently, mice were given an anesthesia antidote consisting of tegmestic (0.3mg/ml; Reckitt Benckiser AG, Switzerland), annexate (0.1mg/ml; Swissmedic, Switzerland) and antisedan (0.1mg/ml; Orion Pharma, Finland) and left to fully recover. Following the surgery, ketoprofen (10mg/kg; Richter Pharma AG, Austria) was subcutaneously injected daily for at least 3 days to reduce any post-operative pain. Animals were given 3-4 weeks to fully recover from the surgery and to allow for expression of the transgene prior to the scanning session.

Another cohort of 13 C57BL/6 mice was injected with 950nl of ssAAV8-mCAMKIIα-hM3D-mCherry of a physical titer ≥ 1.8 x 10^12^ vg/ml following identical procedures as outlined above.

### Electrophysiological recordings

The electrophysiological data used for verification of neuronal activity were acquired in 4 wt-hM3Dq animals. Briefly, animals were anesthetized with isoflurane to match conditions used in imaging studies (2-4% for induction, 0.5% for data collection), and their body temperature was maintained using a heating pad. A small craniotomy was performed over the right and left somatosensory cortex and the brain was covered with silicone oil. A small trepanation was performed over the cerebellum and a silver wire was placed in contact with the CSF to serve as reference electrode. For hM3Dq validation, one silicon probe (Atlas Neurotechnologies, 16 sites, 100 μm spacing) was implanted in each hemisphere. After implantation, we waited 20-30 minutes in order to allow the recording to stabilize. After stabilization, the broadband voltage was amplified and digitally sampled at a rate of 30 kHz or 48 kHz using one of two commercial extracellular recording systems (Intan or Axona). The raw voltage traces were filtered off-line in order to separate the multi-unit activity (bandpass filter 0.46-6 kHz) using a second-order Butterworth filter. Subsequently, the high-passed data were thresholded at 4.5 times the standard deviation across the recording session and the number of spikes in 10 s windows were counted. Recording sessions lasted for 45 min. At 15 min following the start of the recordings (baseline) 30 μg/kg clozapine was injected (intravenously). In order to combine data across mice, the activity at sites with clear multi-unit activity was expressed in percent of the baseline value, i.e., each 10 s bin of MUA activity was divided by the average spike rate during the 15 min pre-injection baseline (100%). All multi-units were then combined from the injected or control region.

### MRI setup and animal preparation

Two MRI sessions were performed on a 7T Bruker BioSpec scanner equipped with a Pharmascan magnet, with two coil setups optimized for the two different acquisition sequences i.e. Arterial Spin Labeling (ASL) and resting-state fMRI (rsfMRI). First, the ASL was obtained utilizing a receiver-only surface coil, coupled with a volume resonator for radiofrequency (rf) transmission. For rsfMRI measurements, a high signal-to-noise ratio (SNR) receive-only cryogenic coil (Bruker BioSpin AG, Fällanden, Switzerland) was used in combination with a linearly polarized room temperature volume resonator for rf transmission.

Standardized anesthesia protocols and animal monitoring procedures were utilized for both MRI sessions^4^. Briefly, mice were initially anesthetized with 4% isoflurane in 1:4 O_2_ to air mixture for 4 min to allow for endotracheal intubation and tail vein cannulation. Mice were positioned on an MRI-compatible support, equipped with hot water-flowing bed to keep the temperature of the animal constant throughout the entire measurement (36.6 ± 0.5°C). The animals were fixed with ear bars and mechanically ventilated via a small animal ventilator (CWE, Ardmore, USA) at the rate of 80 breaths per minute, with 1.8 ml/min flow with isoflurane at 2%. Subsequently, a bolus containing a mixture of medetomidine (0.05 mg/kg) and pancronium (0.25 mg/kg) was injected via the cannulated vein and isoflurane lowered at 1%. Five minutes following the bolus injection, a continuous infusion of medetomidine (0.1 mg/kg/h) and pancronium (0.25 mg/kg/h) was started while isoflurane was further reduced to 0.5%. Animal preparation took on average 15.5 ± 2.7 minutes and all animals fully recovered within 10 min after the measurement.

### Arterial Spin Labeling (ASL)

Brain perfusion was measured using an established arterial spin labelling (ASL) method using a flow sensitive alternating inversion recovery (FAIR) sequence^56, 57^. Briefly, a two-segment Spin-Echo was used with following parameters: echo time TE=12.47 ms, recovery time TR=13000 ms, image matrix=128 x 96, field of view=20 x 20 mm, slice thickness=1 mm, spatial resolution=0.156 x 0.208 x 1 mm/pixel. Inversion parameters: inversion time=40 ms, inversion slab thickness=4 mm, slice margin = 1.5 mm. Sixteen images at different inversion times (50 ms to 3 s) were obtained for T_1_ calculations, resulting in a scan time of 15 min (referred to as ‘baseline’). After the first baseline acquisition, clozapine was intravenously injected at 10-30 μg/kg and three more sessions were acquired (i.e., Post 1, Post 2 and Post 3, respectively), resulting in a total scan time of one hour. A total of 26 animals (5 controls, 8 wt-hM3Dq and 13 CAMK-hM3Dq mice) were scanned.

### Resting-state fMRI (rsfMRI)

Acquisition parameters were following: repetition time TR=1s, echo time TE=15ms, flip angle= 60°, matrix size = 90×50, in-plane resolution = 0.2×0.2 mm^2^, number of slices = 20, slice thickness = 0.4 mm, 3600 volumes for a total scan of 60 min. Clozapine was intravenously injected 15 min after the scan start at the doses of 10 μg/kg or 30 μg/kg. A total of 40 C57BL/6 animals (13 controls and 14 wt-hM3Dq and 13 CAMK-hM3Dq mice) were scanned. These include 8 C57BL/6 animals that were scanned twice (4 controls and 4 wt-hM3Dq mice), once with 10 μg/kg dose of clozapine and another with a 30 μg/kg dose of clozapine (Table S1).

#### Data preprocessing

Data was preprocessed using an already established pipeline for removal of artefacts from the time-series^28, 58^. Briefly, each 4D dataset was normalized in a study-specific EPI template (Advanced Normalization Tools, ANTs v2.1, picsl.upenn.edu/ANTS) and fed into MELODIC (Multivariate Exploratory Linear Optimized Decomposition of Independent Components) to perform within subject special-ICA with a fixed dimensionality estimation (number of components set to 60). The procedure included motion correction and in-plane smoothing with a 0.3 mm kernel. FSL-FIX study-specific classifier, obtained from an independent dataset of 15 mice, was used to perform a ‘conservative’ removal of the variance of the artefactual components^59^. Subsequently, the dataset was despiked, band-pass filtered (0.01-0.25 Hz) and finally normalized into AMBMC template (www.imaging.org.au /AMBMC) using ANTs. Each dataset was split into four parts of 900 data points (equivalent of 15 min of scanning). The difference between the baseline (first 15 min of scan) and the rest of the bins are further referred to as Δ1, Δ2 and Δ3.

#### Seed to seed analysis

The “somatosensory network” in the mouse was defined anatomically using the tracer-based axonal projection pattern of SSctx from the Allen Mouse Brain Connectivity atlas (experiment no. 114290938) and contained contralateral primary SSctx, bilateral somatomotor cortex (MOctx), bilateral association cortices (TEa), as well as ipsilateral caudoputamen (CP) and thalamus (TH (Parafascicular nucleus))^60^. BOLD time series from these regions, apart from thalamus, were extracted using a voxel cube of 3 mm. Thalamus was excluded because previous study from our lab has shown a rather limited FC of cortico-thalamic pathways^60^ detected by rsfMRI. Correlation Z-scored matrices were calculated using FSLNets (FMRIB Analysis Group).

#### Regional Homogeneity (ReHo)

Voxel-wise Regional Homogeneity (ReHo) maps were computed using AFNI. ReHo was calculated for a given voxel and its 19 nearest neighbors and smoothened by Gaussian kernel with Full Width of Half Maximum (FWHM) equal to 10.

#### Voxel-mirrored homotopic connectivity (VMHC)

VMHC measures the similarity between any pair of symmetric inter-hemispheric voxels by computing the Pearson’s correlation coefficient between the time series of each voxel and that of its exact symmetrical inter-hemispheric counterpart. VMHC was computed for all the four timeseries bins and normalized to the baseline for each mouse.

#### Functional connectome analysis

Whole-brain correlation matrices were obtained from a parcellation scheme of 130 regions using the Allen Mouse Brain ontology^23^. In total, 65 regions of interest (ROIs) per hemisphere were considered, including regions from isocortex, hippocampal formation, cortical subplate, striato-pallidum, thalamus, midbrain and hindbrain. Cohen’s d effect size was calculated, using the time series obtained from these whole-brain correlation matrices.

#### Statistical analysis

Prior to performing statistical tests, within-subject normalization to the baseline was implemented. FSL General Linear Model (GLM) was used to perform statistical comparison between controls and wt-hM3Dq mice for the rs-fMRI data (including seed to seed analysis, ReHo, VMHC, connectome analysis). We performed nonparametric permutation testing with 5000 permutations, using family-wise error correction with threshold-free cluster enhancement (TFCE). Statistical significance was defined as *p* < 0.05. CBF differences were statistically tested using linear mixed models implemented in SPSS24 (IBM, USA). All the statistical analysis included 6 covariates: (i) mouse age at the time of surgery; (ii) bodyweight at the time of the scan; (iii) total scan preparation time i.e., time between the first anaesthesia induction to the start of the resting-state scan; (iv) bolus time i.e., time from the first anaesthesia induction to the start of the continuous anaesthetic infusion (isoflurane at 0.5%); (v) the number of ICA components rejected using FSL-FIX and (vi) clozapine dose. These covariates were implemented in all the statistical tests for all types of analysis performed.

### Procedures for PVCre (hM4Di) mice study

#### Transfection procedures for PVCre-hM4Di mice

The right primary somatosensory cortex of PVCre mice (mice expressing Cre recombinase in parvalbumin-expressing neurons) was unilaterally targeted with 950 nl of ssAAV8-hSyn1-dlox-hM4Di_mCherry(rev)-dlox-WPRE-hGHp(A) (n = 19) of a physical titer ≥ 5.4 x 10^12^ vg/ml (*vector genomes; fluorometric quantification) at the rate of 0.06 μL/min. The same procedures for animal preparation, anaesthesia and coordinates were used as already described for wildtype mice.

### In vivo electrophysiology

Four PVCre-hM4Di mice underwent in vivo electrophysiological recordings following the same procedures as described for wildtype mice (Section 2.1.2). The only difference was in the location of the control probe (Atlas Neurotechnologies, 32 sites, 100 μm spacing), which was implanted into the right (ipsilateral to the DREADD injection site) striatum.

### RsfMRI acquisition

Preparation and anesthesia used for the scanning of the PVCre-hM4Di mice was identical to the procedures outlined for wildtype mice (Section 2.1.3). The two acquisitions only differed in the length of the scan. The PVCre-hM4Di mice were scanned for 45 min where clozapine was intravenously injected at 15 min at the dose of 30 mg/ml.

### Classification of univariate BOLD time series

Note that all code for analysis of univariate BOLD dynamics can be found at https://github.com/benfulcher/hctsa_DREADD.

### Data processing and feature computation

We analyzed univariate BOLD time series measured from three brain regions: (i) right SSctx (injected region); (ii) left SSctx (contralateral region); and (iii) visual cortex (control region). Time-series measurements were split into three 15 min durations and labeled according to three classes of experiment: (i) wt-hM3Dq (14 mice); (ii) PVCre-hM4Di (19 mice); and (iii) sham controls (13 mice). To understand which time-series properties distinguish different experimental conditions, we converted each univariate time series to a vector of 7873 interpretable properties (or features) using v0.9.6 of the *hctsa* toolbox^26, 27^. In each brain region, measured time-series data were then represented as a 108 (experiments) *×* 7873 (time-series features) data matrix. Features with well-behaved, real-numbered outputs across the whole dataset and were non-constant within all the three groups were retained for further analysis (7279 features). Each time series was labeled by its experimental condition (‘wt-hM3Dq’, ‘PVCre-hM4Di’, or ‘sham’) and time-point (‘baseline’, ‘t2’, and ‘t3’).

### Classification

We used the feature-based representations of BOLD time-series in each brain area as the basis for classifying the different experimental conditions. We focused our analysis on the Δ1 time period, first subtracting time-series features computed at baseline, and then normalizing these feature differences using an outlier-robust sigmoidal transformation^27^. A linear support vector machine classification model was learned on the normalized feature matrix for a given brain area, using inverse probability class reweighting to accommodate class imbalance. A measure of discriminability of a pair of classes was quantified as the balanced classification accuracy using 10-fold stratified cross validation. Balanced accuracy was computed as the arithmetic mean of sensitivity and specificity to account for the small class imbalance: 13 sham controls and 14 wt-hM3Dq or 19 PVCre-hM4Di mice. To reduce variance in the random partition of data into 10 folds, we repeated this procedure 50 times (with each iteration yielding a balanced accuracy value), and summarized the resulting distribution of balanced accuracies as its mean and standard deviation.

In smaller samples, there is a greater probability that optimistic classification results can be obtained by chance. To quantify this, we evaluated the statistical significance of our classification results relative to random assignments of class labels to data. This was achieved by computing a null distribution of the same performance metric used above (mean across 50 repeats of 10-fold crossvalidated balanced accuracy) for 5000 random class label assignments to time series. The statistical significance of a given classification result was then estimated as a permutation test (as the proportion of 5000 null samples with a mean balanced classification rate exceeding that of the true assignment of class labels).

### Low-dimensional projections

As explained above, each BOLD signal was represented as a feature vector, or equivalently, as a point in a high-dimensional feature space. To aid visualization of the class structure of data in this space, we projected normalized time series × feature data matrices into a lower-dimensional space that captures the maximal variance in the full feature space using principal components analysis. In these plots, individual time series are points in the space, providing an intuitive visualization of the structure of the time-series dataset.

### Interpreting differences in univariate dynamics

We aimed to determine which properties of the univariate BOLD dynamics in the injected brain area best discriminated: (i) wt-hM3Dq from controls; (ii) PVCre-hM4Di from controls; and (iii) wt-hM3Dq from PVCre-hM4Di mice. Focusing on the first time period after baseline, we first computed the difference in each feature value between this first time period and the baseline period (Δ1). We then tested whether these relative feature values differed between groups using a Wilcoxon rank-sum test. Statistical correction for the large number of hypothesis tests was performed as the false discovery rate^61^, setting a significance threshold of *q* = 0.05.

### Predicting *Fmr1^y/-^* mice using a classifier trained on PVCre-hM4Di

Time series from 46 cortical ROIs (23 in each hemisphere) obtained from *Fmr1^y/-^* (*n* = 44) and wildtype littermates (n = 39) was obtained from two of our previous studies^4, 53^. The classification proceeded in two stages. In the training phase, a classification model was trained on normalized features of rs-fMRI time series in SSctx (right hemisphere) during the Δ1 time period, as described above. That is, the trained classifier learns features of the BOLD dynamics to accurately predict whether a new time series is PVCre-hM4Di or control. In the testing phase, the trained model was used to predict labels of rsfMRI time series in *Fmr1^y/-^* dataset. We identified PVCre-hM4Di with the *Fmr1^y/-^* knockout condition, and quantified prediction performance as balanced accuracy (%) as above, evaluated in each brain region independently. P-values were estimated for each brain area using permutation testing with 5 × 10^6^ null samples generated by independent shuffling of labels, and corrected across brain regions (focusing just on the right hemisphere) using the false discovery rate method of Benjimini and Hochberg^61^.

### Histological evaluation of transfection

DREADDs viral expression (for both wt-hM3Dq and PVCre-hM4Di) was confirmed by m-Cherry staining using standard immunohistochemistry protocols. Briefly, after the last MRI session mice were deeply anesthetized using a mixture of Ketamine (100mg/kg; Graeub, Switzerland), Xylazine (10mg/kg; Rompun, Bayer) and Acepromazine (2mg/kg; Fatro S.p.A, Italy) and transcardially perfused with 4% Paraformaldehyde (PFA, pH=7.4). The brains were postfixed in 4% PFA for 1.5 hours at 4°C and then placed overnight in 30% sucrose solution. Brains were frozen in a tissue mounting fluid (Tissue-Tek O.C.T Compund, Sakura Finetek Europe B.V., Netherlands) and sectioned coronally in 40 μm thick slices using a cryostat (MICROM HM 560, histocom AG-Switzerland). Free-floating slices were first permeabilized in 0.2% Triton X-100 for 30 min and then incubated overnight in 0.2% Triton X-100, 2% normal goat serum and rabbit anti-mCherry (1:1000, Ab167453, Abcam) at 4°C under continuous agitation (100rpm/min). The next day sections were incubated for 1h in 0.2% Triton X-100, 2% normal goat serum, goat anti-rabbit Alexa Flour 546 (1:300, A11035, Life Technologies) and Nissl (1:300, NeuroTrace 660, Thermo Molecular Probes) or DAPI (1:300, Sigma-Aldrich) at room temperature under continuous agitation. Afterwards, slices were mounted on the superfrost slides where they were left to airdry and later coverslipped with Dako Flourescence mounting medium (Agilent Technologies). Confocal laser-scanning microscope (Leica, SP8, Centre for Microscopy and Image Analysis, UZH) and Zeiss Slidescanner (Zeiss Axio scan, Z1, Centre for Microscopy and Image Analysis, UZH) were used to detect the viral expression. Microscopy protocol included a tile scan with a 10x or a 20x objective, pixel size of 1.2μm and image size of 1024×1024 pixels. Images were preprocessed and analyzed using ImageJ-Fiji and Zeiss Zen Blue, respectively.

## Abbreviations

ASD: autism spectrum disorder
ASL: arterial spin labelling
BOLD: blood-oxygen level dependent
CAMKII: alpha-calcium/calmodulin-dependent protein kinase II promoter
CBF: cerebral blood flow
CP: caudate putamen
DREADD: Designer Receptors Exclusively Activated by Designer Drugs
FC: functional connectivity
Fmr1: fragile X mental retardation 1 gene
hM3Dq: excitatory DREADD
hM4Di: inhibitory DREADD
MOctx: primary somatomotor cortex
PV: parvalbumin interneurons
PVCre: parvalbumin cre-dependant
PVCre-hM4Di: PVCre mice injected with hM4Di DREADD
ReHo: Regional Homogeneity
rsfMRI: resting-state functional Magnetic Resonance Imaging
SSctx: primary somatosensory cortex
STR: striatum
TEa: temporal association cortex
TH: thalamus
VMHC: Voxel-mirrored Homotopic Connectivity
Wt-hM3Dq: wildtype mice injected with hM3Dq DREADD

## 6. Acknowledgment

We thank the team of the EPIC animal facility for providing animal care. We thank Jean-Charles Paterna from the Viral Vector Facility (VVF) of the Neuroscience Center Zurich, a joint competence center of ETH Zurich and University of Zurich for producing viral vectors and viral vector plasmids. We thank Amalia Floriou Servou and Mattia Privitera for useful feedback regarding immunohistochemistry.

## 8. Author Contributions

M.M. performed viral injections, conducted the fMRI experiments, analyzed the data, performed immunohistochemistry and wrote the paper. B.D.F. performed machine-learning analysis and wrote the paper. C.L. conducted electrophysiology experiments, analyzed the data and wrote the paper (results, method). F.H. supervised electrophysiology and provided feedback on the study. M.R. supervised fMRI experiments and provided feedback on the study. V.Z and N.W. designed the study, supervised fMRI experiments, analyzed data and wrote the paper.

