## Supplementary Files for "Cortical excitation:inhibition imbalance causes abnormal brain network dynamics as observed in neurodevelopmental disorders"

**Supplementary Figures**

***Figure S1. Schematic of the experimental pipeline.***

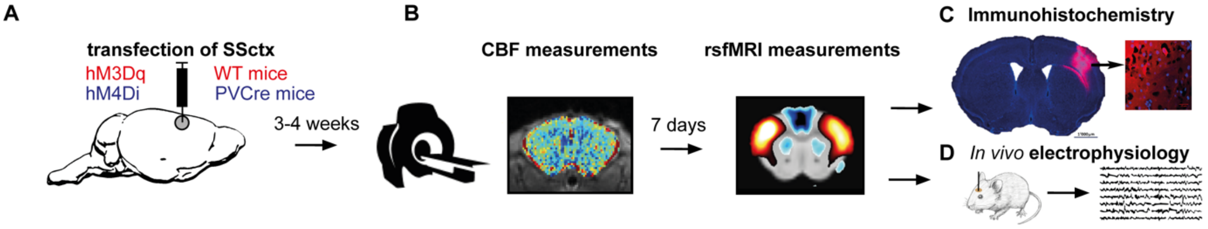


***Figure S1. A)*** *Administration of excitatory (hM3Dq) or inhibitory (hM4Di) DREADD constructs via viral injection into the right primary somatosensory cortex (SSctx) of wildtype and PVCre mice, respectively.* ***B)*** *Three to four weeks after surgery, mice underwent multimodal MRI functional imaging for the assessment of cerebral blood flow and resting-state fMRI (rsfMRI). In each session, clozapine was injected i.v. (<=0.03 mg/kg) to activate the DREADD during the recordings.* ***C)*** *After the rsfMRI session, the successful viral transfection was ensured by immunohistochemistry as depicted by the results from a single brain slice.* ***D)*** *In some mice,* in vivo *electrophysiological recordings were taken from the right SSctx and from a control region, to confirm that DREADD-transfected neurons are successfully and selectively activated after clozapine injection and cause a net shift towards over-excitation.*

***Figure S2. hM3Dq activated by CNO causes inter-hemispheric connectivity reduction***

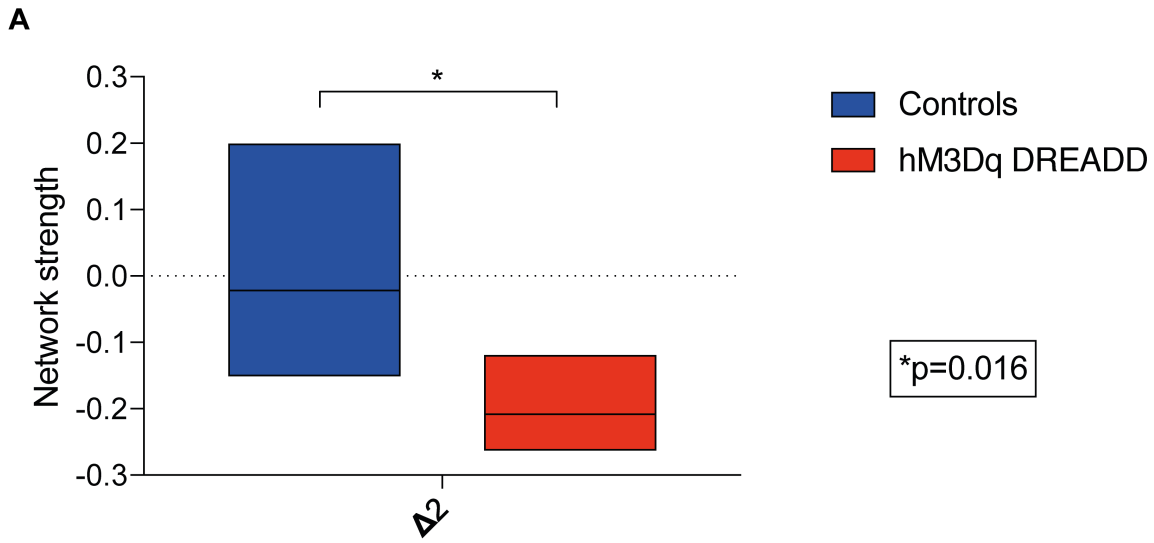


***Figure S2.*** *Graph represents a decrease in interhemispheric connectivity between the somatomotor regions in wt-hM3Dq mice (n = 6) compared to controls (n = 5), occurring 15 min after the CNO i.v. injection. The DREADD was activated using 1 mg/kg CNO. Univariate analysis of ANOVA show a significant difference between the groups F(1,9) = 8.710, p = 0.02.*

*Figure S3. Short- and long-range connectivity changes induced by activating hM3Dq*

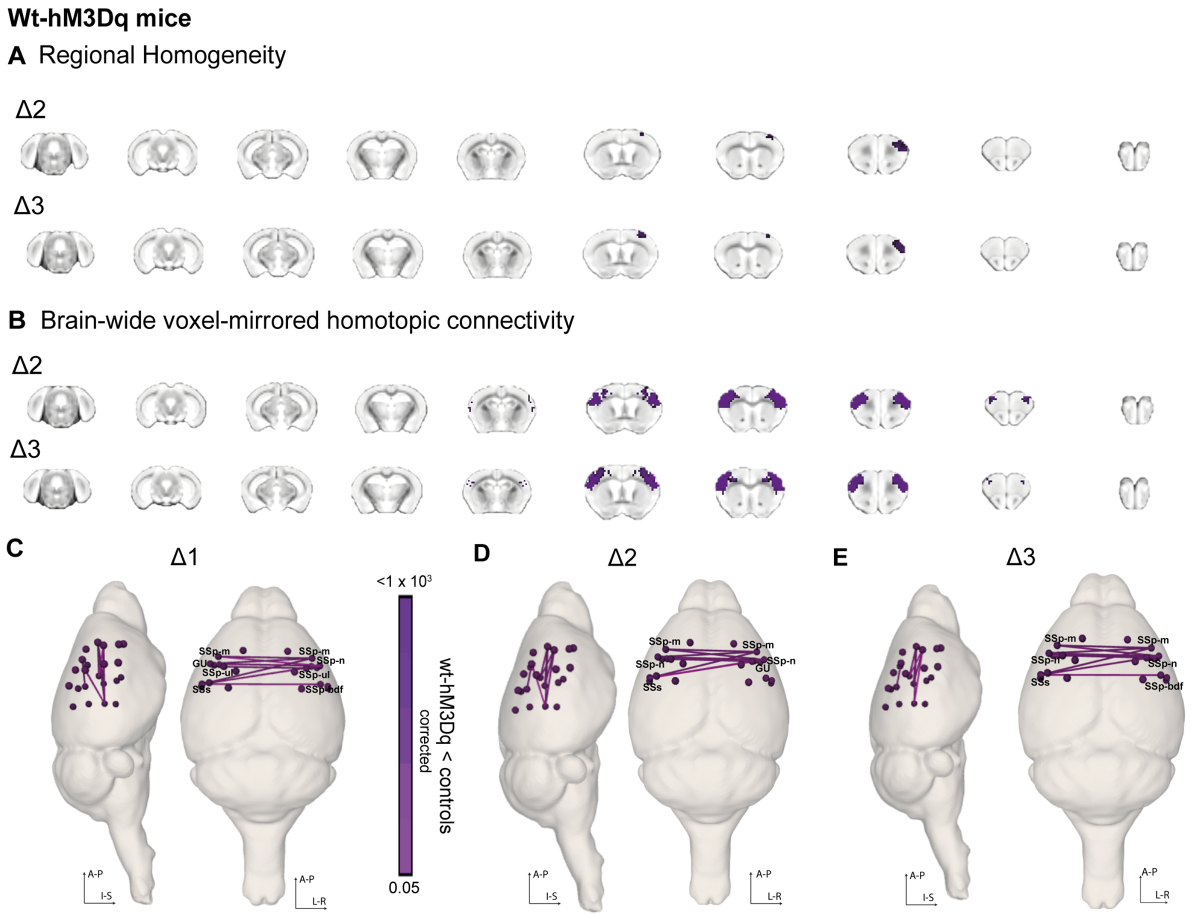


***Figure S3. A)*** *Significant decrease (corrected) in regional homogeneity shown through coronal slices for Δ2 and Δ3 time periods.* ***B)*** *Significant (corrected) interhemispheric decrease in VMHC shown through coronal slices for Δ2 and Δ3 time periods.* ***C)*** *Whole-brain connectome analysis shows a significant interhemispheric reduction between somatosensory cortices for Δ1, Δ2 and Δ3 time periods between wt-hM3Dq (n=14) mice and controls (n=13). Regions affected are as follows: SSp-m: Primary Somatosensory Area, mouth; GU: Gustatory areas; SSp-ul: Primary Somatosensory Area, upper limb; SSs: Supplementary somatosensory area**;** SSp-bdf: Primary Somatosensory Area, barrel field; SSp-n: Primary Somatosensory Area, nose**.***

*Figure S4. Long-range connectivity changes induced by activating hM3Dq*

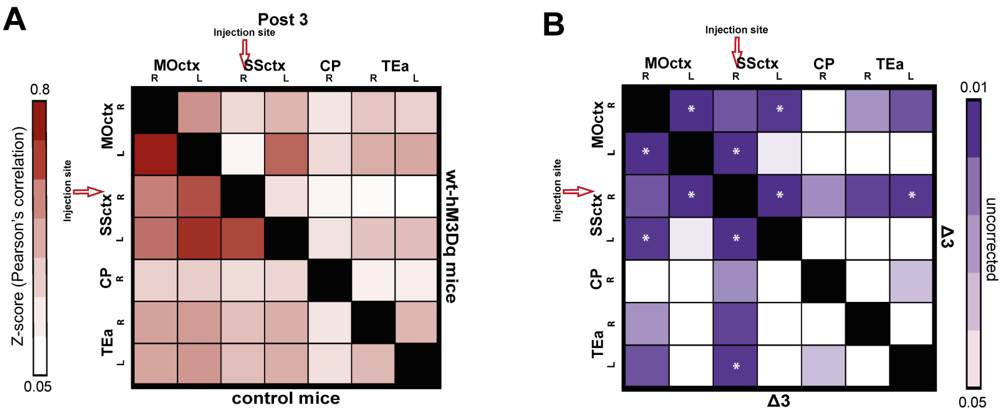


***Figure S4. A)** Averaged and z-scored Pearson's correlation of all control (n=13) animals (lower triangle of the matrix) and all wt-hM3Dq (n=14) animals (upper triangle of the matrix) for Post 3. **B)** Statistically significant differences during Δ3 time period among 4 brain regions (MOctx – Somatomotor cortex, SSctx – somatosensory cortex, CP – caudoputamen and TEa – temporal association cortex, R – right, L - left) between the controls and wt-hM3Dq mice. The stars within some ROIs indicate statistically significant differences that survived correction of multiple comparisons.*

***Figure S5. Changes induced by activating CAMKII-hM3Dq in wildtype mice.***

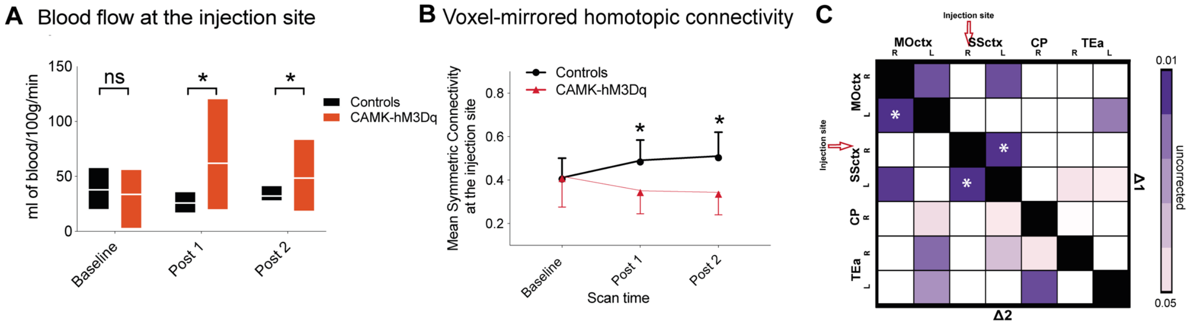


***Figure S5. A)*** *Comparison of blood flow (ml of blood/100g/min) over time between controls and wildtype mice injected with excitatory (hM3Di) DREADD on CAMKII promoter (measured at the injection site). We observe a significant (repeated measures ANOVA indicates a groups*scan time effect: F(1.0,20.0)=7.3;p=0.004) increase in blood flow in CAMK-hM3Dq mice (n=13) after clozapine injection (similar to wt-hM3Dq from Fig. 2H).* ***B)*** *Change over time of mean symmetric connectivity at the injection site. There is a significant decrease (repeated measures ANOVA shows a groups*scan time effect: F(1.16, 21.98)=8.21, p=0.007) in mean connectivity in CAMK-hM3Dq (n=13) mice as compared to controls (n=13).* ***C)*** *Matrix represents statistically significant differences during Δ1 and Δ2 time period among 4 brain regions (MOctx – Somatomotor cortex, SSctx – somatosensory cortex, CP –caudoputamen and TEa – temporal association cortex, R – right, L - left) between the controls and CAMK-hM3Dq mice. The stars within some ROIs indicate statistically significant differences that survived correction of multiple comparisons.*

***Figure S6. Changes induced by activating hM4Di in PVCre mice.***

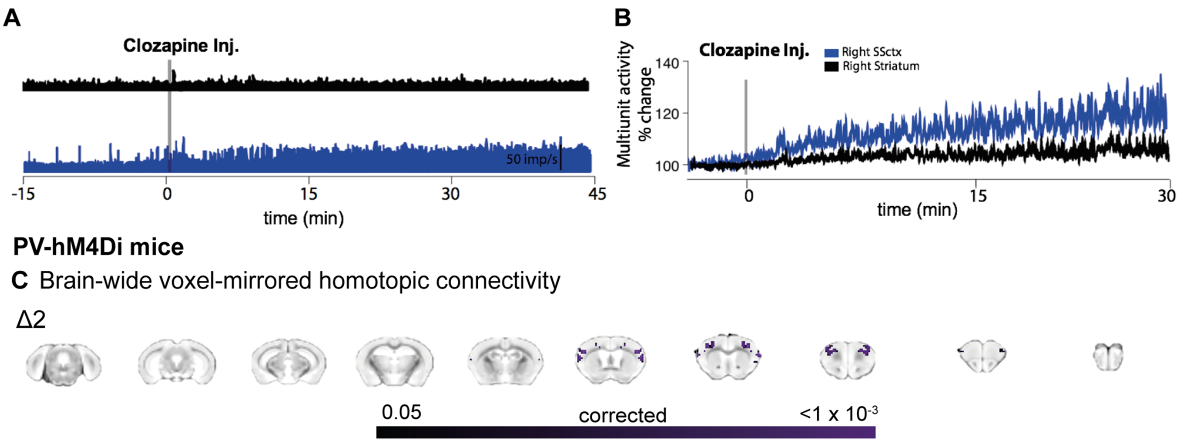


***Figure S6. A)*** *Time-resolved firing rate in millisecond bins for the right (blue) somatosensory cortex and right striatum (black). A steady increase of firing rate occurs at the right SSctx once clozapine is injected, while no change occurs in striatum.* ***B)*** *Averaged multiunit activity increase by almost 20% compared to baseline (before clozapine injection) in right SSctx and remained similar compared to baseline in the striatum.* ***C)*** *Significant decrease (TFCE-corrected) in symmetric connectivity in PVCre-hM4Di mice compared to controls depicted on coronal slices for the Δ2 time point.*

***Figure S7. Viral expressions in the SSctx of wt-hM3Dq, PVCre-hM4Di and CAMKII-hM3Dq mice***

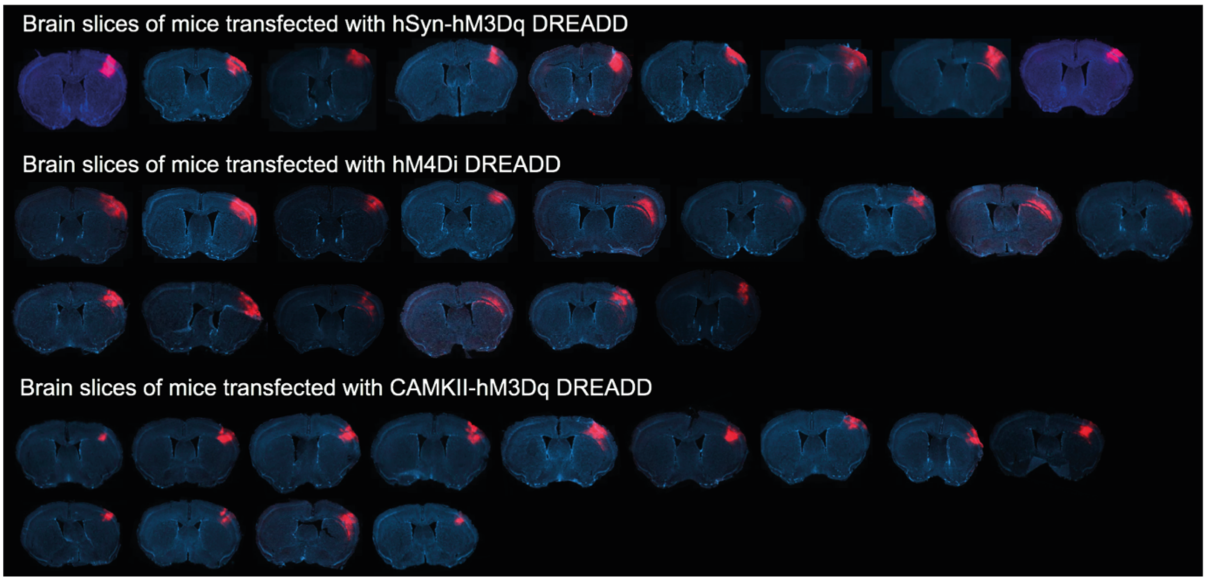


***Figure S7.*** *Viral expressions at the somatosensory cortex of the wt-hSyn-hM3Dq DREADD, PVCre-hM4Di and for wt-CAMKII-hM3Dq mice. Each slice represents the expression obtained from a mouse.*

***Figure S8. Statistical comparison between different doses of clozapine in wt-hM3Dq and control mice***

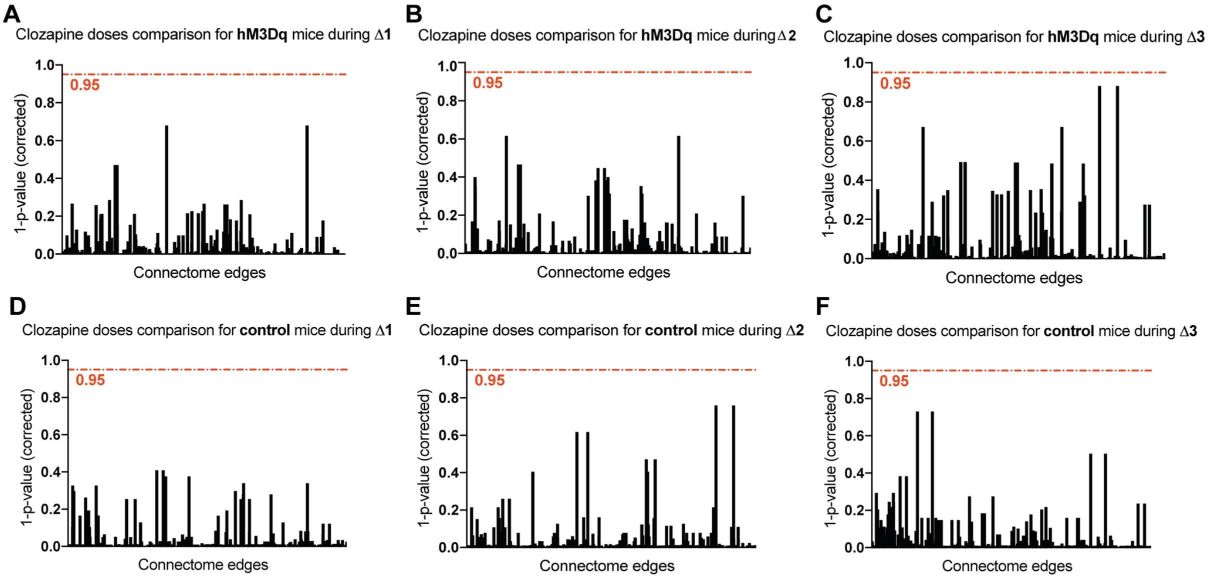


***Figure S8. A,B,C)*** *Corrected statistical comparison (1-p-value) between the control groups that received 2 different doses of clozapine (10 mg/kg or 30 mg/kg) for the Δ1, Δ2 and Δ3 time periods. Red line represents a start of significance, showing that there are no significant differences between the groups.* ***D,E,F)*** *Similar information to the 3 graphs above but only for the mice transfected with hM3Dq DREADD.*

***Figure S9. Univariate BOLD dynamics change in similar ways in wt-hM3Dq and PVCre-hM4Di***

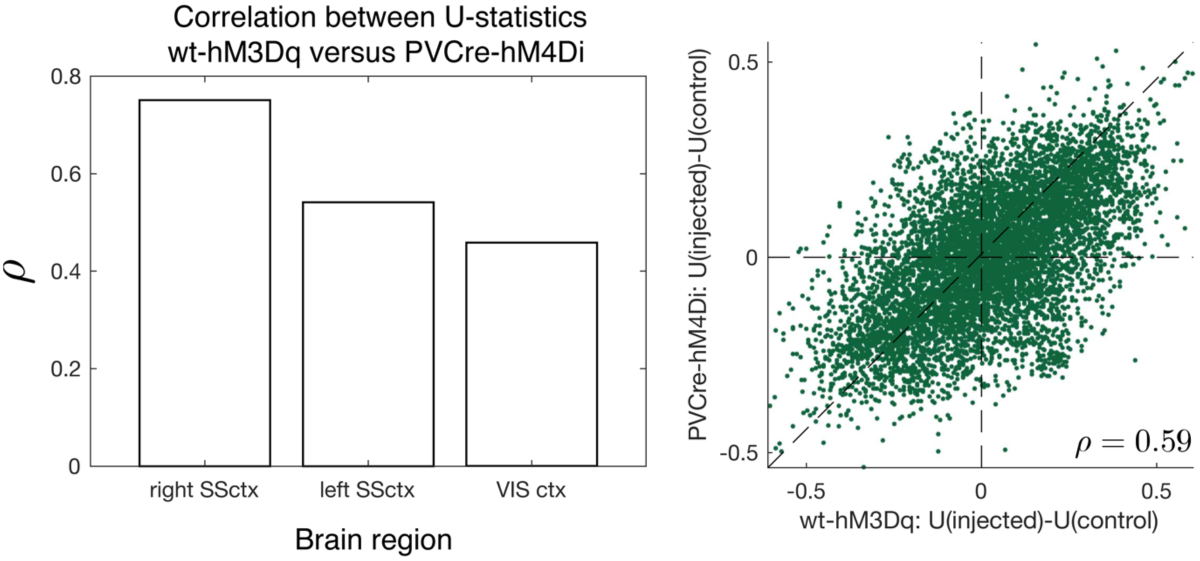


***Figure S9. A)*** *We measured similarity of dynamical changes in wt-hM3Dq and PVCre-hM4Di relative to control as the correlation, $\rho$, of Mann-Whitney U statistics computed for each feature at $\Delta 1$, which are plotted as bars for each brain region.* ***B)*** *After subtracting matched test statistics measured in VISctx from those measured in right SSctx, the residual correlation in the injected region remains strong, shown here as a scatter plot.*

***Table S1. Demographics of mice used in the experiment***

| Mouse ID | Genotype | Age at surgery (days) | Bodyweight at surgery (gr) | Type of DREADD | Gender | Bodyweight on the scan day (gr) | Clozapine Dose (μg/kg) |
|---|---|---|---|---|---|---|---|
| 86 | C57BL/6 | 84 | 23.4 | excitatory | male | 25.1 | 10 |
| 89 | C57BL/6 | 84 | 22.7 | excitatory | male | 24.3 | 10 |
| 90 | C57BL/6 | 84 | 22.9 | excitatory | male | 23.4 | 10 |
| 101 | C57BL/6 | 51 | 21.9 | excitatory | male | 25.1 | 10 |
| 102 | C57BL/6 | 51 | 24.7 | excitatory | male | 25.6 | 10 |
| 103 | C57BL/6 | 51 | 24.9 | excitatory | male | 28.2 | 10 |
| 104 | C57BL/6 | 51 | 25 | excitatory | male | 25.5 | 10 |
| 105 | C57BL/6 | 51 | 27.1 | excitatory | male | 27.6 | 10 |
| 109 | C57BL/6 | 52 | 24.4 | excitatory | male | 25 | 10 |
| 110 | C57BL/6 | 52 | 25.2 | excitatory | male | 27 | 10 |
| 101 | C57BL/6 | 51 | 21.9 | excitatory | male | 26.2 | 30 |
| 103 | C57BL/6 | 51 | 24.9 | excitatory | male | 29.3 | 30 |
| 104 | C57BL/6 | 51 | 25 | excitatory | male | 23.3 | 30 |
| 110 | C57BL/6 | 52 | 25.2 | excitatory | male | 29.1 | 30 |
| | average ⟹ | **58.29 ± 13.94** | **24.23 ± 1.47** | | | **26.05 ± 1.94** | |
| 70 | C57BL/6 | 86 | 30 | sham | male | 28.6 | 10 |
| 72 | C57BL/6 | 86 | 20.8 | sham | male | 23.6 | 10 |
| 73 | C57BL/6 | 86 | 21 | sham | male | 23.3 | 10 |
| 75 | C57BL/6 | 86 | 22 | sham | male | 24 | 10 |
| 119 | C57BL/6 | 53 | 24.9 | sham | male | 25.5 | 10 |
| 121 | C57BL/6 | 54 | 26.6 | sham | male | 27.3 | 10 |
| 123 | C57BL/6 | 54 | 26.2 | sham | male | 26.6 | 10 |
| 124 | C57BL/6 | 54 | 26.7 | sham | male | 28.6 | 10 |
| 125 | C57BL/6 | 54 | 25 | sham | male | 25.9 | 10 |
| 121 | C57BL/6 | 54 | 26.6 | sham | male | 29 | 30 |
| 123 | C57BL/6 | 54 | 26.2 | sham | male | 29.6 | 30 |
| 124 | C57BL/6 | 54 | 26.7 | sham | male | 28.1 | 30 |
| 125 | C57BL/6 | 54 | 25 | sham | male | 27.1 | 30 |
| | average ⟹ | **63.77 ± 15.43** | **23.21 ± 2.6** | | | **26.71 ± 2.12** | |
| 8 | PVCre | 89 | 22 | inhibitory | female | 26.5 | 30 |
| 9 | PVCre | 89 | 20.3 | inhibitory | female | 20.7 | 30 |
| 10 | PVCre | 89 | 21.1 | inhibitory | female | 22.5 | 30 |
| 11 | PVCre | 81 | 28.6 | inhibitory | male | 29.3 | 30 |
| 12 | PVCre | 81 | 30.7 | inhibitory | male | 33 | 30 |
| 13 | PVCre | 81 | 27.3 | inhibitory | male | 28.1 | 30 |
| 14 | PVCre | 89 | 22 | inhibitory | female | 24.7 | 30 |
| 5 | PVCre | 89 | 28.9 | inhibitory | male | 29.5 | 30 |
| 6 | PVCre | 90 | 29.6 | inhibitory | male | 26.1 | 30 |
| 7 | PVCre | 89 | 19.8 | inhibitory | female | 21.7 | 30 |
| 100 | PVCre | 61 | 24.5 | inhibitory | male | 25.1 | 30 |
| 64 | PVCre | 98 | 25.2 | inhibitory | male | 25.8 | 30 |
| 65 | PVCre | 98 | 27.2 | inhibitory | male | 28 | 30 |
| 71 | PVCre | 82 | 30 | inhibitory | male | 29.3 | 30 |
| 76 | PVCre | 82 | 26.9 | inhibitory | male | 27.8 | 30 |
| 87 | PVCre | 75 | 27.9 | inhibitory | male | 28.2 | 30 |
| 89 | PVCre | 75 | 27.5 | inhibitory | male | 27.9 | 30 |
| 92 | PVCre | 61 | 27.5 | inhibitory | male | 30.1 | 30 |
| 93 | PVCre | 61 | 27.7 | inhibitory | male | 29.7 | 30 |
| | average ⟹ | **82.11 ± 11.3** | **26.04 ± 3.4** | | | **27.1 ± 1.40** | |
| 185 | C57BL/6 | 66 | 23.8 | CAMK - excitatory | male | 27.3 | 30 |
| 188 | C57BL/6 | 66 | 24.2 | CAMK - excitatory | male | 26 | 30 |
| 190 | C57BL/6 | 67 | 24.5 | CAMK - excitatory | male | 25.7 | 30 |
| 195 | C57BL/6 | 68 | 26.7 | CAMK - excitatory | male | 26 | 30 |
| 199 | C57BL/6 | 67 | 23.8 | CAMK - excitatory | male | 27.6 | 30 |
| 192 | C57BL/6 | 67 | 22.3 | CAMK - excitatory | male | 25.8 | 30 |
| 193 | C57BL/6 | 67 | 24.1 | CAMK - excitatory | male | 26.6 | 30 |
| 194 | C57BL/6 | 67 | 25.8 | CAMK - excitatory | male | 28.2 | 30 |
| 200 | C57BL/6 | 76 | 26.6 | CAMK - excitatory | male | 29.3 | 30 |
| 201 | C57BL/6 | 76 | 27 | CAMK - excitatory | male | 29.2 | 30 |
| 202 | C57BL/6 | 76 | 25.6 | CAMK - excitatory | male | 27.2 | 30 |
| 203 | C57BL/6 | 76 | 23 | CAMK - excitatory | male | 24.7 | 30 |
| 204 | C57BL/6 | 76 | 26.1 | CAMK - excitatory | male | 28.4 | 30 |
| | average ⟹ | **70.4 ± 4.6** | **24.9 ± 1.5** | | | **27.1 ± 1.4** | |

***Table S1.*** *Details about all the mice involved in the experiments with mean + standard deviation calculated for mouse age at surgery, bodyweight at surgery and bodyweight on the scan day.*

### Supplementary Results and Discussion

#### Activated DREADDs caused relative over-excitation at the cellular level and within cortical circuits

In our first experiment, we used hM3Dq-DREADD on human synapsin (hSyn) promoter to achieve a strong shift in E:I balance towards over-excitation in the targeted region. According to a study from Nathanson and colleagues, the hSyn promoter in AAV constructs is able to generate a clear bias towards gene expression in excitatory cortical neurons (up to 85%), which is similar to results with CAMKII of similar titer [1]. To address this, we injected a cohort of mice with a hM3Dq-DREADD on a CAMKII promoter and confirmed similar results as with hSyn, thus showing that both hM3Dq DREADD on a CAMKII or hSyn promoter generate identical results (Fig S6). While none of the promoters in AAV can restrict its affinity exclusively toward excitatory neurons, our electrophysiological results confirmed that activating DREADDs in the wt-hM3Dq mice caused a significant increase of neuronal firing rate in the targeted region, suggesting a general over-excitation. Convergent evidence is also revealed by our CBF measurements in both wt-hM3Dq and CAMK-hM3Dq mice. Numerous PET and fMRI studies have demonstrated that CBF reflects glucose metabolism, and it is considered an indirect marker of neural activity at the population level [2, 3]. Here we report an increase in CBF exclusively in the right SSctx of DREADD mice, which confirms the anatomical specificity of our intervention in exciting the targeted area.

In the second set of experiments, we used PVCre mice to ensure cell-specificity of the hM4Di DREADD. Analogous to our first experiment, *in vivo* electrophysiology confirmed that inhibiting GABAergic PV neurons via hM4Di significantly increased neuronal population firing and moderately increased blood flow. This finding is consistent with previous literature, which states that suppression of PV interneurons leads to an increased cortical excitation i.e. increase in number of cells spiking at the population level [4, 5]. In summary, even though the two experiments targeted different biological pathways, both manipulations shifted cortical E:I balance in a similar manner by causing a net increase in the firing rate of neuronal populations within the targeted SSctx, as shown by convergent evidence from *in vivo* electrophysiology and cerebral blood flow.

In order to activate DREADDs, we used low-doses of clozapine instead of the more commonly used clozapine-N-oxide (CNO). Clozapine is a potent antipsychotic medication used in the treatment of schizophrenia that binds to a number of receptors including serotonin, α1-adrenergic receptors, muscarinic-1 and histamine [6]. Clozapine is a metabolite of CNO but, unlike CNO, is able to freely pass the blood brain barrier. Because of its high affinity for DREADD receptors, clozapine is thought to be the actual activator of DREADDs in *in vivo* studies [7, 8]. In our experiments, we used clozapine doses of 0.01–0.03 mg/kg to activate the DREADDs, which are below the threshold level necessary to elicit a behavioral effect in mice. To further control for any unspecific effect of clozapine, we employed a randomized controlled design injecting identical doses of clozapine in sham-operated mice. We also conducted experiments using two different doses of clozapine (Fig. S7) and found no differences in any of the groups, suggesting that the lower dose (0.01 mg/kg) is sufficient to activate DREADDs *in vivo*. Furthermore, we performed analogous experiments in smaller mouse cohorts using CNO, which revealed similar effects, i.e., a significant decrease in interhemispheric connectivity in the targeted somatomotor cortex (Fig. S3).

Mildly anesthetized animals were used during the experiment in order to minimize animal movement and potential distress during examination. Increasing body of literature emphasizes that the importance for understanding the effects of anesthesia on functional connectivity is key to correct interpretation of the results [9, 10, 11]. For optimal results, we used a combination of medetomidine (vasoconstrictor), isoflorane (vasodilator) and pancronium (muscle relaxant) at very low doses [11, 12]. Isoflorane is predominantly targeting the GABAergic neurotransmitter system [13]. GABA receptors are widely spread throughout the brain, therefore it is essential to control for percentage of isoflurane used in the anesthesia. Metedomidine predominantly targets α2 adrenergic receptors, whose distribution within the brain varies, being largely located in the brainstem (especially locus coeruleus) [13]. By combining metedomidine with isoflurane we ensured that isoflurane is kept at minimum level (0.5%) and as so does not have a dominant effect on functional connectivity.

**Univariate BOLD dynamics change in similar ways in both wt-hM3Dq and PVCre-hM4Di**

We aimed to test whether changes in univariate BOLD dynamics resulting from both DREADDs mechanisms (wt-hM3Dq and PVCre-hM4Di) relative to controls were similar, focusing on the Δ1 time period. For each time-series feature, we quantified how the DREADDs condition differed from the sham condition as a Mann-Whitney U statistic, and then compared these statistics between the two different DREADDs. If the properties of BOLD dynamics change similarly due to wt-hM3Dq and PVCre-hM4Di DREADDs, then we should detect this as an increase in correlation, $\rho$, between the two sets of Mann-Whitney U statistics. Indeed, we found that this correlation, $\rho$, is lowest in the control region (VISctx, $\rho = 0.46$), increases in the contralateral region (left SSctx, $\rho = 0.54$), and is highest in the injected region (right SSctx, $\rho = 0.75$), as shown in Fig. S6A. The two sets of Mann-Whitney U statistics (wt-hM3Dq versus control and PVCre-hM4Di versus control) are not independent due to a repeated comparison to the same set of control dynamics, likely accounting for the correlation observed in the VISctx. After subtracting matched test statistics between the right SSctx and VISctx, correlations between the residuals of this subtraction are strong in both the right SSctx ($\rho_{\text{resid}} = 0.59$, shown in Fig. S8B), and left SSctx ($\rho_{\text{resid}} = 0.47$).

**Discriminatory Time-Series Features**

Having demonstrated that wt-hM3Dq and PVCre-hM4Di manipulations cause distinctive changes to local BOLD dynamics, particularly in the injected region, we aimed to understand the specific properties of BOLD dynamics that drive this discriminability. We first investigated which time-series features most strongly differentiate wt-hM3Dq from controls in the right SSctx at Δ1. We scored the discriminability of each feature separately using a rank-sum test, correcting for multiple-hypothesis testing across all individual features by assuming independent tests [14] (a conservative correction due to high inter-correlation of many time-series features in *hctsa* [15]). A total of 924 features of wt-hM3Dq BOLD dynamics were significantly different to control ($p_{corr} < 0.05$), including measures of variance, which were decreased in wt-hM3Dq mice (e.g., standard deviation, $p_{corr} = 2 \times 10^{-3}$), and measures of autocorrelation, which were increased in wt-hM3Dq mice (e.g., power in the lowest fifth of frequencies, $p_{corr} = 3 \times 10^{-3}$). A wide range of other features from diverse literatures also captured a significant difference in BOLD dynamics in wt-hM3Dq mice relative to controls, including features of self-affine scaling, linear and nonlinear autocorrelation, local motifs, temporal entropy, model-based fits and forecasting, wavelet decompositions, outliers, and stationarity. All features with $p_{corr} < 0.01$ are grouped broadly by conceptual theme and listed in Supplementary File 1. Our results indicate a robust time-series signature of wt-hM3Dq activation.

We repeated the above analysis to identify whether we can isolate informative changes to BOLD dynamics caused by PVCre-hM4Di. We identified 225 time-series features that are individually informative of the difference from controls ($p_{corr} < 0.05$). Significant features are dominated by various measures of autocorrelation in the signal, similar to the characteristics identified above for wt-hM3Dq. Features are listed in Supplementary File 2.

**BOLD time-series classification at Δ2 and Δ3**

Our main analysis of classifying univariate BOLD dynamics was focused on the Δ1 time period. Here we demonstrate that the main qualitative results reported hold also for later time periods Δ2 (and Δ3, available only for wt-hM3Dq and sham). Namely: the control region (VISctx) was consistently classified at chance level ($p > 0.05$), the contralateral region (left SSctx) mostly displayed a classifiability intermediate between the control and injected regions, while the injected region (right SSctx) consistently displayed the highest classification rates. Individual results are listed here:

- wt-hM3Dq versus CONTROL at Δ2: right SSctx: 86% ($p = 2 \times 10^{4}$), left SSctx: 38% ($p = 0.8$), VISctx: 53% ($p = 0.4$).
- wt-hM3Dq versus CONTROL at Δ3: right SSctx: 75% ($p = 0.01$), left SSctx : 49% ($p = 0.5$), VISctx: 45% ($p = 0.6$).
- PVCre-hM4Di versus CONTROL at Δ2: right SSctx: 66% ($p = 0.05$), left SSctx: 72% ($p = 0.01$), VISctx: 52% ($p = 0.4$).
- wt-hM3Dq versus PVCre-hM4Di at Δ2: right SSctx: 68% ($p = 0.03$), left SSctx: 52% ($p = 0.4$), VISctx: 36% ($p = 0.9$).

Here, we list features of **wt-hM3Dq mice** from hctsa (v0.96) with corrected p < 0.01

Features have been coarsely grouped into categories, as labeled.

Each row is of the form:

featureID | featureName | featureKeywords | pValue | pValueCorr

---Distribution---
7872 | MD_rawHRVmeas_SD1 | medical,raw,spreaddep | 6.98e-07 | 0.00135
33 | DN_Moments_raw_4 | distribution,moment,shape,raw,spreaddep | 2.99e-06 | 0.00193
22 | median_absolute_deviation | distribution,spread,raw,spreaddep | 9.67e-06 | 0.00202
35 | DN_Moments_raw_6 | distribution,moment,shape,raw,spreaddep | 4.49e-06 | 0.00202
16 | rms | distribution,location,raw,locdep,spreaddep | 1.39e-05 | 0.00221
19 | standard_deviation | distribution,spread,raw,spreaddep | 1.39e-05 | 0.00221
20 | mean_absolute_deviation | distribution,spread,raw,spreaddep | 1.94e-05 | 0.00221
37 | DN_Moments_raw_8 | distribution,moment,shape,raw,spreaddep | 1.94e-05 | 0.00221
7873 | MD_rawHRVmeas_SD2 | medical,raw,spreaddep | 2.71e-05 | 0.00247
21 | interquartile_range | distribution,spread,raw,spreaddep | 3.72e-05 | 0.00271
930 | DN_FitKernelSmoothraw_entropy | distribution,ksdensity,entropy,raw,spreaddep | 5.06e-05 | 0.00308
39 | DN_Moments_raw_10 | distribution,moment,shape,raw,spreaddep | 6.78e-05 | 0.00369

---Powerlaw scaling---
4120 | SC_FluctAnal_2_dfa_50_2_logi_r2_se1 | scaling | 2.99e-06 | 0.00193
3961 | SC_FluctAnal_2_std_50_logi_ssr | scaling | 9.67e-06 | 0.00202
4062 | SC_FluctAnal_2_dfa_50_0_logi_meanssr | scaling | 9.67e-06 | 0.00202
750 | SC_fastdfa_exponent | dfa,scaling,mex | 1.94e-05 | 0.00221
3974 | SC_FluctAnal_2_std_50_logi_r2_linfitint | scaling | 1.94e-05 | 0.00221
4057 | SC_FluctAnal_2_dfa_50_0_logi_ssr | scaling | 1.94e-05 | 0.00221
4070 | SC_FluctAnal_2_dfa_50_0_logi_r2_linfitint | scaling | 2.71e-05 | 0.00247
4089 | SC_FluctAnal_2_dfa_50_1_logi_r1_alpha | scaling | 2.71e-05 | 0.00247
3966 | SC_FluctAnal_2_std_50_logi_meanssr | scaling | 3.72e-05 | 0.00271
4103 | SC_FluctAnal_2_dfa_50_2_logi_se1 | scaling | 6.78e-05 | 0.00369
4104 | SC_FluctAnal_2_dfa_50_2_logi_se2 | scaling | 6.78e-05 | 0.00369
4079 | SC_FluctAnal_2_dfa_50_1_logi_se1 | scaling | 9.02e-05 | 0.00394
4080 | SC_FluctAnal_2_dfa_50_1_logi_se2 | scaling | 9.02e-05 | 0.00394

4121 | SC_FluctAnal_2_dfa_50_2_logi_r2_se2 | scaling | 9.02e-05 | 0.00394

4145 | SC_FluctAnal_2_dfa_50_3_logi_r2_se2 | scaling | 9.02e-05 | 0.00394

4133 | SC_FluctAnal_2_dfa_50_3_logi_ratsplitminerr | scaling | 0.000119 | 0.00455

3902 | SC_FluctAnal_2_nothing_50_logi_r2_linfitint | scaling | 0.000155 | 0.00541

3985 | SC_FluctAnal_2_iqr_50_logi_ssr | scaling | 0.000155 | 0.00541

4016 | SC_FluctAnal_2_rsrange_50_logi_r1_linfitint | scaling | 0.000155 | 0.00541

4144 | SC_FluctAnal_2_dfa_50_3_logi_r2_se1 | scaling | 0.000155 | 0.00541

4017 | SC_FluctAnal_2_rsrange_50_logi_r1_alpha | scaling | 0.000201 | 0.00617

4037 | SC_FluctAnal_2_rsrangefit_50_1_logi_ratsplitminerr | scaling | 0.000201 | 0.00617

4088 | SC_FluctAnal_2_dfa_50_1_logi_r1_linfitint | scaling | 0.000201 | 0.00617

3885 | SC_FluctAnal_2_nothing_50_logi_linfitint | scaling | 0.000259 | 0.0071

4055 | SC_FluctAnal_2_dfa_50_0_logi_se1 | scaling | 0.000259 | 0.0071

4056 | SC_FluctAnal_2_dfa_50_0_logi_se2 | scaling | 0.000259 | 0.0071

4182 | SC_FluctAnal_2_dfa_50_1_3_logi_meanssr | scaling | 0.000259 | 0.0071

4033 | SC_FluctAnal_2_rsrangefit_50_1_logi_ssr | scaling | 0.00033 | 0.00819

4086 | SC_FluctAnal_2_dfa_50_1_logi_meanssr | scaling | 0.00033 | 0.00819

4153 | SC_FluctAnal_2_dfa_50_1_2_logi_ssr | scaling | 0.00033 | 0.00819

3937 | SC_FluctAnal_2_range_50_logi_ssr | scaling | 0.000419 | 0.00934

3948 | SC_FluctAnal_2_range_50_logi_r1_ssr | scaling | 0.000419 | 0.00934

3950 | SC_FluctAnal_2_range_50_logi_r2_linfitint | scaling | 0.000419 | 0.00934

4163 | SC_FluctAnal_2_dfa_50_1_2_logi_r1_se2 | scaling | 0.000419 | 0.00934

---Fourier Power Spectrum---

4312 | SP_Summaries_pgram_hamm_fpolysat_r2 | spectral | 9.67e-06 | 0.00202

4436 | SP_Summaries_welch_rect_fpolysat_r2 | spectral | 9.67e-06 | 0.00202

4437 | SP_Summaries_welch_rect_fpolysat_rmse | spectral | 6.68e-06 | 0.00202

4442 | SP_Summaries_welch_rect_ylogareatopeak | spectral | 9.67e-06 | 0.00202

4493 | SP_Summaries_welch_rect_logarea_5_1 | spectral | 4.49e-06 | 0.00202

4560 | SP_Summaries_fft_fpolysat_r2 | spectral | 9.67e-06 | 0.00202

4561 | SP_Summaries_fft_fpolysat_rmse | spectral | 6.68e-06 | 0.00202

4566 | SP_Summaries_fft_ylogareatopeak | spectral | 9.67e-06 | 0.00202

4617 | SP_Summaries_fft_logarea_5_1 | spectral | 4.49e-06 | 0.00202

4430 | SP_Summaries_welch_rect_fpoly2csS_p3 | spectral | 1.94e-05 | 0.00221

4483 | SP_Summaries_welch_rect_logarea_4_1 | spectral | 1.94e-05 | 0.00221

4554 | SP_Summaries_fft_fpoly2csS_p3 | spectral | 1.94e-05 | 0.00221

4607 | SP_Summaries_fft_logarea_4_1 | spectral | 1.94e-05 | 0.00221

4313 | SP_Summaries_pgram_hamm_fpolysat_rmse | spectral | 1.89e-06 | 0.00183

4431 | SP_Summaries_welch_rect_fpoly2_sse | spectral | 2.71e-05 | 0.00247

4433 | SP_Summaries_welch_rect_fpoly2_rmse | spectral | 2.71e-05 | 0.00247

4555 | SP_Summaries_fft_fpoly2_sse | spectral | 2.71e-05 | 0.00247

4557 | SP_Summaries_fft_fpoly2_rmse | spectral | 2.71e-05 | 0.00247

4494 | SP_Summaries_welch_rect_area_5_2 | spectral | 3.72e-05 | 0.00271

4618 | SP_Summaries_fft_area_5_2 | spectral | 3.72e-05 | 0.00271

4307 | SP_Summaries_pgram_hamm_fpoly2_sse | spectral | 6.78e-05 | 0.00369

4399 | SP_Summaries_welch_rect_w_weighted_peak_height | spectral | 6.78e-05 | 0.00369

4523 | SP_Summaries_fft_w_weighted_peak_height | spectral | 6.78e-05 | 0.00369

4360 | SP_Summaries_pgram_hamm_area_4_2 | spectral | 9.02e-05 | 0.00394

4398 | SP_Summaries_welch_rect_w_weighted_peak_prom | spectral | 9.02e-05 | 0.00394

4432 | SP_Summaries_welch_rect_fpoly2_r2 | spectral | 9.02e-05 | 0.00394

4434 | SP_Summaries_welch_rect_fpolysat_a | spectral | 9.02e-05 | 0.00394

4492 | SP_Summaries_welch_rect_area_5_1 | spectral | 9.02e-05 | 0.00394

4495 | SP_Summaries_welch_rect_logarea_5_2 | spectral | 9.02e-05 | 0.00394

4522 | SP_Summaries_fft_w_weighted_peak_prom | spectral | 9.02e-05 | 0.00394

4556 | SP_Summaries_fft_fpoly2_r2 | spectral | 9.02e-05 | 0.00394

4558 | SP_Summaries_fft_fpolysat_a | spectral | 9.02e-05 | 0.00394

4616 | SP_Summaries_fft_area_5_1 | spectral | 9.02e-05 | 0.00394

4619 | SP_Summaries_fft_logarea_5_2 | spectral | 9.02e-05 | 0.00394

4417 | SP_Summaries_welch_rect_tau | spectral | 9.65e-05 | 0.00413

4541 | SP_Summaries_fft_tau | spectral | 9.65e-05 | 0.00413

4306 | SP_Summaries_pgram_hamm_fpoly2csS_p3 | spectral | 0.000119 | 0.00455

4309 | SP_Summaries_pgram_hamm_fpoly2_rmse | spectral | 0.000119 | 0.00455

4420 | SP_Summaries_welch_rect_wmax_25 | spectral | 0.000112 | 0.00455

4435 | SP_Summaries_welch_rect_fpolysat_b | spectral | 0.000119 | 0.00455

4544 | SP_Summaries_fft_wmax_25 | spectral | 0.000112 | 0.00455

4559 | SP_Summaries_fft_fpolysat_b | spectral | 0.000119 | 0.00455

4458 | SP_Summaries_welch_rect_linfitloglog_mf_a1 | spectral | 0.000155 | 0.00541

4582 | SP_Summaries_fft_linfitloglog_mf_a1 | spectral | 0.000155 | 0.00541

4370 | SP_Summaries_pgram_hamm_area_5_2 | spectral | 0.000201 | 0.00617

4427 | SP_Summaries_welch_rect_w25_75 | spectral | 0.000191 | 0.00617

4551 | SP_Summaries_fft_w25_75 | spectral | 0.000191 | 0.00617

4297 | SP_Summaries_pgram_hamm_centroid | spectral | 0.000235 | 0.00709

4296 | SP_Summaries_pgram_hamm_wmax_25 | spectral | 0.000259 | 0.0071

4419 | SP_Summaries_welch_rect_wmax_10 | spectral | 0.000296 | 0.008

4543 | SP_Summaries_fft_wmax_10 | spectral | 0.000296 | 0.008

4429 | SP_Summaries_welch_rect_fpoly2csS_p2 | spectral | 0.00033 | 0.00819

4553 | SP_Summaries_fft_fpoly2csS_p2 | spectral | 0.00033 | 0.00819

4421 | SP_Summaries_welch_rect_centroid | spectral | 0.00035 | 0.00859

4545 | SP_Summaries_fft_centroid | spectral | 0.00035 | 0.00859

4482 | SP_Summaries_welch_rect_area_4_1 | spectral | 0.000419 | 0.00934

4484 | SP_Summaries_welch_rect_area_4_2 | spectral | 0.000419 | 0.00934

4606 | SP_Summaries_fft_area_4_1 | spectral | 0.000419 | 0.00934

4608 | SP_Summaries_fft_area_4_2 | spectral | 0.000419 | 0.00934

4732 | SP_Summaries_fft_logdev_area_4_2 | spectral | 0.000419 | 0.00934

7762 | MD_hrv_classic_hf | medical | 0.000201 | 0.00617

---Linear autocorrelation---

6944 | NL_embed_PCA_mi_10_std | embedding,pca | 1.94e-05 | 0.00221

95 | AC_3 | correlation | 5.06e-05 | 0.00308

6958 | NL_embed_PCA_1_10_perc_2 | embedding,pca | 6.78e-05 | 0.00369

6940 | NL_embed_PCA_mi_10_perc_2 | embedding,pca | 0.000119 | 0.00455

6959 | NL_embed_PCA_1_10_perc_3 | embedding,pca | 0.000119 | 0.00455

6948 | NL_embed_PCA_mi_10_nto50 | embedding,pca | 0.000128 | 0.00489

6947 | NL_embed_PCA_mi_10_top2 | embedding,pca | 0.000155 | 0.00541

6949 | NL_embed_PCA_mi_10_nto60 | embedding,pca | 0.000171 | 0.00591

6950 | NL_embed_PCA_mi_10_nto70 | embedding,pca | 0.0002 | 0.00617

6945 | NL_embed_PCA_mi_10_range | embedding,pca | 0.000259 | 0.0071

6965 | NL_embed_PCA_1_10_top2 | embedding,pca | 0.00033 | 0.00819

6939 | NL_embed_PCA_mi_10_perc_1 | embedding,pca | 0.000419 | 0.00934

94 | AC_2 | correlation | 0.000259 | 0.0071

---Nonlinear autocorrelation properties---

244 | CO_HistogramAMI_even_5_3 | information,correlation,AMI | 2.99e-06 | 0.00193

194 | RM_ami_3 | information,correlation,AMI | 4.49e-06 | 0.00202

154 | AC_nl_33 | correlation,nonlinearautocorr | 9.67e-06 | 0.00202

177 | AC_nl_025 | correlation,nonlinearautocorr | 1.94e-05 | 0.00221

224 | CO_HistogramAMI_std2_2_3 | information,correlation,AMI | 6.68e-06 | 0.00202

249 | CO_HistogramAMI_even_10_3 | information,correlation,AMI | 6.68e-06 | 0.00202

493 | CO_glscf_1_1_3 | correlation,glscf | 9.67e-06 | 0.00202

234 | CO_HistogramAMI_std2_10_3 | information,correlation,AMI | 1.39e-05 | 0.00221

209 | CO_HistogramAMI_std1_2_3 | information,correlation,AMI | 1.39e-05 | 0.00221

214 | CO_HistogramAMI_std1_5_3 | information,correlation,AMI | 1.94e-05 | 0.00221

229 | CO_HistogramAMI_std2_5_3 | information,correlation,AMI | 1.94e-05 | 0.00221
5812 | SD_TSTL_surrogates_1_100_2_tc3_stdsurr | nonlinear,tstool,correlation,surrogate | 1.94e-05 | 0.00221
324 | IN_AutoMutualInfoStats_40_kraskov1_4_ami3 | information,correlation,AMI | 1.94e-05 | 0.00221
219 | CO_HistogramAMI_std1_10_3 | information,correlation,AMI | 2.71e-05 | 0.00247
239 | CO_HistogramAMI_even_2_3 | information,correlation,AMI | 2.71e-05 | 0.00247
499 | CO_glscf_1_2_3 | correlation,glscf | 2.71e-05 | 0.00247
517 | CO_glscf_2_2_3 | correlation,glscf | 2.71e-05 | 0.00247
166 | AC_nl_003 | correlation,nonlinearautocorr | 3.72e-05 | 0.00271
259 | CO_HistogramAMI_quantiles_5_3 | information,correlation,AMI | 3.72e-05 | 0.00271
521 | CO_glscf_2_5_1 | correlation,glscf | 3.72e-05 | 0.00271
165 | AC_nl_002 | correlation,nonlinearautocorr | 5.06e-05 | 0.00308
264 | CO_HistogramAMI_quantiles_10_3 | information,correlation,AMI | 5.06e-05 | 0.00308
269 | IN_AutoMutualInfoStats_40_gaussian_ami3 | information,correlation,AMI | 5.06e-05 | 0.00308
1133 | CO_trev_3_denom | correlation,nonlinear | 5.06e-05 | 0.00308
474 | CO_CompareMinAMI_even_2_80_std | correlation,AMI | 5.14e-05 | 0.0031
187 | AC_nl_122 | correlation,nonlinearautocorr | 6.78e-05 | 0.00369
1108 | CO_tc3_3_denom | correlation,nonlinear | 6.78e-05 | 0.00369
470 | CO_CompareMinAMI_even_2_80_max | correlation,AMI | 8.32e-05 | 0.00394
139 | AC_nl_12345 | correlation,nonlinearautocorr | 9.02e-05 | 0.00394
160 | AC_nl_033 | correlation,nonlinearautocorr | 9.02e-05 | 0.00394
170 | AC_nl_012 | correlation,nonlinearautocorr | 9.02e-05 | 0.00394
182 | AC_nl_036 | correlation,nonlinearautocorr | 9.02e-05 | 0.00394
185 | AC_nl_112 | correlation,nonlinearautocorr | 9.02e-05 | 0.00394
189 | AC_nl_223 | correlation,nonlinearautocorr | 9.02e-05 | 0.00394
254 | CO_HistogramAMI_quantiles_2_3 | information,correlation,AMI | 9.02e-05 | 0.00394
181 | AC_nl_026 | correlation,nonlinearautocorr | 0.000119 | 0.00455
186 | AC_nl_113 | correlation,nonlinearautocorr | 0.000119 | 0.00455
190 | AC_nl_233 | correlation,nonlinearautocorr | 0.000119 | 0.00455
471 | CO_CompareMinAMI_even_2_80_range | correlation,AMI | 0.000108 | 0.00455
171 | AC_nl_013 | correlation,nonlinearautocorr | 0.000155 | 0.00541
515 | CO_glscf_2_2_1 | correlation,glscf | 0.000155 | 0.00541
1128 | CO_trev_2_denom | correlation,nonlinear | 0.000201 | 0.00617
1264 | CO_AddNoise_1_gaussian_ami_at_5 | correlation,AMI,entropy | 0.000201 | 0.00617
509 | CO_glscf_1_10_1 | correlation,glscf | 0.000259 | 0.0071
1273 | CO_AddNoise_1_gaussian_fitlinb | correlation,AMI,entropy | 0.000259 | 0.0071

477 | CO_CompareMinAMI_even_2_80_modef | correlation,AMI | 0.000303 | 0.00812
1198 | CO_AddNoise_1_even_10_fitlinb | correlation,AMI,entropy | 0.00033 | 0.00819
1189 | CO_AddNoise_1_even_10_ami_at_5 | correlation,AMI,entropy | 0.000419 | 0.00934
490 | CO_CompareMinAMI_std2_2_80_nlocmax | correlation,AMI | 0.000425 | 0.00943

---Symbolic Motifs and Surprise---
1406 | SB_TransitionMatrix_3ac_sumdiagcov | symbolic,transitionmat | 1.39e-05 | 0.00221
3517 | SB_MotifTwo_diff_ddud | symbolic,motifs | 1.21e-05 | 0.00221
3642 | SB_MotifThree_quantile_abaa | symbolic,motifs | 1.84e-05 | 0.00221
1411 | SB_TransitionMatrix_3ac_maxeigcov | symbolic,transitionmat | 3.72e-05 | 0.00271
1413 | SB_TransitionMatrix_3ac_stdeigcov | symbolic,transitionmat | 3.72e-05 | 0.00271
3636 | SB_MotifThree_quantile_aaba | symbolic,motifs | 3.59e-05 | 0.00271
1417 | SB_TransitionMatrix_4ac_TD4 | symbolic,transitionmat | 4.86e-05 | 0.00308
3508 | SB_MotifTwo_diff_dud | symbolic,motifs | 4.47e-05 | 0.00308
3608 | SB_MotifThree_quantile_aba | symbolic,motifs | 4.3e-05 | 0.00308
2474 | FC_Surprise_T1_10_1_m2quad_500_median | information,symbolic | 5.72e-05 | 0.00343
1401 | SB_TransitionMatrix_3ac_T9 | symbolic,transitionmat | 7.31e-05 | 0.00394
1452 | SB_TransitionpAlphabet_40_1_maxdiagfexp_b | symbolic,transitionmat | 9.02e-05 | 0.00394
3519 | SB_MotifTwo_diff_dudd | symbolic,motifs | 8.23e-05 | 0.00394
3768 | SB_MotifThree_diffquant_abca | symbolic,motifs | 9.62e-05 | 0.00413
2342 | FC_Surprise_dist_5_2_udq_500_std | information,symbolic | 0.000119 | 0.00455
3560 | SB_MotifTwo_mean_uuud | symbolic,motifs | 0.000165 | 0.00571
3553 | SB_MotifTwo_mean_duuu | symbolic,motifs | 0.000212 | 0.00647
1399 | SB_TransitionMatrix_3ac_T7 | symbolic,transitionmat | 0.000213 | 0.00648
1451 | SB_TransitionpAlphabet_40_1_maxdiagfexp_a | symbolic,transitionmat | 0.000259 | 0.0071
3713 | SB_MotifThree_quantile_cccc | symbolic,motifs | 0.000272 | 0.00745
1084 | MD_polvar_01_3 | medical,symbolic | 0.00035 | 0.00859
2272 | FC_Surprise_T1_20_2_q_500_min | information,symbolic | 0.000425 | 0.00943

---Entropy---
2887 | EN_Randomize_dyndist_ac2fexpa | entropy,slow | 6.68e-06 | 0.00202
2764 | EN_mse_1-10_2_015_diff1_sampen_s1 | entropy,sampen,controlen,mse | 1.39e-05 | 0.00221
2830 | EN_Randomize_statdist_ac3fexpa | entropy,slow | 1.94e-05 | 0.00221
2893 | EN_Randomize_dyndist_ac3fexpa | entropy,slow | 3.72e-05 | 0.00271
914 | ApEn2_01 | entropy | 5.06e-05 | 0.00308
2956 | EN_Randomize_permute_ac3fexpa | entropy,slow | 5.06e-05 | 0.00308
2746 | EN_SampEn_5_02_diff1_meanchsampen | entropy,sampen,controlen | 0.000201 | 0.00617

913 | ApEn1_02 | entropy | 0.000259 | 0.0071

2905 | EN_Randomize_dyndist_sampen2_015fexpa | entropy,slow | 0.00033 | 0.00819

2824 | EN_Randomize_statdist_ac2fexpa | entropy,slow | 0.000419 | 0.00934

---Model Fit or Model-Based Prediction/Forecasting---

7014 | MF_steps_ahead_ar_2_6_rmserr_4 | model,prediction | 6.98e-07 | 0.00135

7015 | MF_steps_ahead_ar_2_6_mabserr_4 | model,prediction | 3.99e-07 | 0.00135

7006 | MF_steps_ahead_ar_2_6_mabserr_1 | model,prediction | 1.89e-06 | 0.00183

7012 | MF_steps_ahead_ar_2_6_mabserr_3 | model,prediction | 1.89e-06 | 0.00183

7269 | MF_StateSpace_n4sid_1_05_1_acsnd0 | model | 2.99e-06 | 0.00193

3840 | MF_AR_arcov_2_a3 | model,fit | 9.67e-06 | 0.00202

7005 | MF_steps_ahead_ar_2_6_rmserr_1 | model,prediction | 9.67e-06 | 0.00202

7011 | MF_steps_ahead_ar_2_6_rmserr_3 | model,prediction | 6.68e-06 | 0.00202

7025 | MF_steps_ahead_ar_2_6_maxdiffrms | model,prediction | 6.68e-06 | 0.00202

7084 | MF_steps_ahead_arma_3_1_6_mabserr_4 | model,prediction | 4.49e-06 | 0.00202

3020 | FC_LocalSimple_mean4_stderr | forecasting | 1.94e-05 | 0.00221

3041 | FC_LoopLocalSimple_mean_stderr_peaksize | forecasting | 1.94e-05 | 0.00221

7026 | MF_steps_ahead_ar_2_6_stddiffrms | model,prediction | 1.94e-05 | 0.00221

7095 | MF_steps_ahead_arma_3_1_6_stddiffrms | model,prediction | 1.94e-05 | 0.00221

7370 | MF_CompareAR_1_10_all_maxdiff | model | 1.94e-05 | 0.00221

7009 | MF_steps_ahead_ar_2_6_mabserr_2 | model,prediction | 2.71e-05 | 0.00247

7463 | MF_armax_2_2_05_1_AR_2 | model | 2.71e-05 | 0.00247

7129 | MF_FitSubsegments_ar_2_uniform_25_01_a_2_mean | model,prediction | 3.72e-05 | 0.00271

7264 | MF_StateSpace_n4sid_1_05_1_ac3 | model | 3.72e-05 | 0.00271

7267 | MF_StateSpace_n4sid_1_05_1_ac3n | model | 3.72e-05 | 0.00271

7385 | MF_CompareAR_1_10_05_stddiff | model | 3.72e-05 | 0.00271

7386 | MF_CompareAR_1_10_05_maxdiff | model | 3.72e-05 | 0.00271

7415 | MF_armax_3_1_05_1_ac3 | model | 3.72e-05 | 0.00271

7418 | MF_armax_3_1_05_1_ac3n | model | 3.72e-05 | 0.00271

7081 | MF_steps_ahead_arma_3_1_6_mabserr_3 | model,prediction | 5.06e-05 | 0.00308

7088 | MF_steps_ahead_arma_3_1_6_ac1_5 | model,prediction | 5.06e-05 | 0.00308

7258 | MF_StateSpace_n4sid_1_05_1_p2_5 | model | 5.06e-05 | 0.00308

7369 | MF_CompareAR_1_10_all_stddiff | model | 5.06e-05 | 0.00308

3019 | FC_LocalSimple_mean4_meanabserr | forecasting | 6.78e-05 | 0.00369

3070 | FC_LocalSimple_median5_meanabserr | forecasting | 6.78e-05 | 0.00369

7083 | MF_steps_ahead_arma_3_1_6_rmserr_4 | model,prediction | 6.78e-05 | 0.00369

3010 | FC_LocalSimple_mean3_stderr | forecasting | 9.02e-05 | 0.00394

7018 | MF_steps_ahead_ar_2_6_mabserr_5 | model,prediction | 9.02e-05 | 0.00394
7138 | MF_FitSubsegments_arma_2_2_uniform_25_01_fpe_mean | model,prediction | 9.02e-05 | 0.00394
7257 | MF_StateSpace_n4sid_1_05_1_p1_5 | model | 9.02e-05 | 0.00394
7008 | MF_steps_ahead_ar_2_6_rmserr_2 | model,prediction | 0.000119 | 0.00455
7045 | MF_steps_ahead_ar_best_6_ac1_6 | model,prediction,arfit | 0.000119 | 0.00455
7085 | MF_steps_ahead_arma_3_1_6_ac1_4 | model,prediction | 0.000119 | 0.00455
7147 | MF_FitSubsegments_arma_2_2_uniform_25_01_p_2_mean | model,prediction | 0.000119 | 0.00455
3874 | DN_SimpleFit_sin1_resAC1 | model | 0.000155 | 0.00541
3875 | DN_SimpleFit_sin1_resAC2 | model | 0.000155 | 0.00541
7296 | MF_StateSpace_n4sid_2_05_1_p2_5 | model | 0.000155 | 0.00541
7466 | MF_armax_2_2_05_1_maxda | model | 0.000155 | 0.00541
3030 | FC_LocalSimple_meantau_stderr | forecasting | 0.000201 | 0.00617
3038 | FC_LoopLocalSimple_mean_stderr_chn | forecasting | 0.000201 | 0.00617
3879 | DN_SimpleFit_sin2_resAC2 | model | 0.000201 | 0.00617
7017 | MF_steps_ahead_ar_2_6_rmserr_5 | model,prediction | 0.000201 | 0.00617
7087 | MF_steps_ahead_arma_3_1_6_mabserr_5 | model,prediction | 0.000201 | 0.00617
7341 | MF_StateSpace_n4sid_3_05_1_p2_5 | model | 0.000201 | 0.00617
3000 | FC_LocalSimple_mean2_stderr | forecasting | 0.000259 | 0.0071
3061 | FC_LocalSimple_median3_stderr | forecasting | 0.000259 | 0.0071
7091 | MF_steps_ahead_arma_3_1_6_ac1_6 | model,prediction | 0.000259 | 0.0071
7247 | MF_StateSpace_n4sid_1_05_1_k_1 | model | 0.000259 | 0.0071
7263 | MF_StateSpace_n4sid_1_05_1_ac2 | model | 0.000259 | 0.0071
7282 | MF_StateSpace_n4sid_2_05_1_A_4 | model | 0.000259 | 0.0071
7347 | MF_StateSpace_n4sid_3_05_1_ac3 | model | 0.000259 | 0.0071
7350 | MF_StateSpace_n4sid_3_05_1_ac3n | model | 0.000259 | 0.0071
3071 | FC_LocalSimple_median5_stderr | forecasting | 0.00033 | 0.00819
6767 | MF_arfit_1_8_sbc_meanA | modelfit,arfit | 0.00033 | 0.00819
6769 | MF_arfit_1_8_sbc_sumA | modelfit,arfit | 0.00033 | 0.00819
7040 | MF_steps_ahead_ar_best_6_rmserr_5 | model,prediction,arfit | 0.00033 | 0.00819
7148 | MF_FitSubsegments_arma_2_2_uniform_25_01_p_2_max | model,prediction | 0.00033 | 0.00819
7250 | MF_StateSpace_n4sid_1_05_1_m_fpe | model | 0.00033 | 0.00819
7340 | MF_StateSpace_n4sid_3_05_1_p1_5 | model | 0.00033 | 0.00819
7408 | MF_armax_3_1_05_1_p1_5 | model | 0.00033 | 0.00819
7448 | MF_armax_1_1_05_1_ac3 | model | 0.00033 | 0.00819

7451 | MF_armax_1_1_05_1_ac3n | model | 0.00033 | 0.00819
7453 | MF_armax_1_1_05_1_acsnd0 | model | 0.00033 | 0.00819
3120 | FC_LocalSimple_lfit5_stderr | forecasting | 0.000419 | 0.00934
7043 | MF_steps_ahead_ar_best_6_rmserr_6 | model,prediction,arfit | 0.000419 | 0.00934
7048 | MF_steps_ahead_ar_best_6_maxdiffrms | model,prediction,arfit | 0.000419 | 0.00934
7253 | MF_StateSpace_n4sid_1_05_1_meanabs | model | 0.000419 | 0.00934
7262 | MF_StateSpace_n4sid_1_05_1_ac1 | model | 0.000419 | 0.00934
7265 | MF_StateSpace_n4sid_1_05_1_ac1n | model | 0.000419 | 0.00934
7266 | MF_StateSpace_n4sid_1_05_1_ac2n | model | 0.000419 | 0.00934
7272 | MF_StateSpace_n4sid_1_05_1_dwts | model | 0.000419 | 0.00934
7362 | MF_CompareAR_1_10_all_maxv | model | 0.000419 | 0.00934
7441 | MF_armax_1_1_05_1_p1_5 | model | 0.000419 | 0.00934

---Gaussian Processes---
6462 | MF_GP_hyperparameters_covSEiso_covNoise_1_200_resample_mlikelihood | model,GaussianProcess | 0.000155 | 0.00541
6465 | MF_GP_hyperparameters_covSEiso_covNoise_1_200_resample_std_mu_data | model,GaussianProcess | 0.000201 | 0.00617
6475 | MF_GP_hyperparameters_covSEiso_covPeriodic_covNoise_1_200_resample_mlikelihood | model,GaussianProcess | 0.000201 | 0.00617
6477 | MF_GP_hyperparameters_covSEiso_covPeriodic_covNoise_1_200_resample_mabserr_std | model,GaussianProcess | 0.000201 | 0.00617
6478 | MF_GP_hyperparameters_covSEiso_covPeriodic_covNoise_1_200_resample_std_mu_data | model,GaussianProcess | 0.000201 | 0.00617
6476 | MF_GP_hyperparameters_covSEiso_covPeriodic_covNoise_1_200_resample_rmserr | model,GaussianProcess | 0.000259 | 0.0071
6463 | MF_GP_hyperparameters_covSEiso_covNoise_1_200_resample_rmserr | model,GaussianProcess | 0.000419 | 0.00934
6464 | MF_GP_hyperparameters_covSEiso_covNoise_1_200_resample_mabserr_std | model,GaussianProcess | 0.000419 | 0.00934
6481 | MF_GP_hyperparameters_covSEiso_covPeriodic_covNoise_1_200_resample_minS | model,GaussianProcess | 0.000419 | 0.00934

---Wavelet decomposition---
6586 | WL_cwt_db3_32_meanabsC | wavelet,cwt | 5.06e-05 | 0.00308
6612 | WL_cwt_sym2_32_meanabsC | wavelet,cwt | 5.06e-05 | 0.00308
6587 | WL_cwt_db3_32_medianabsC | wavelet,cwt | 9.02e-05 | 0.00394

6753 | WL_dwtcoeff_sym2_5_stdd_l4 | wavelet,dwt | 0.000119 | 0.00455
6613 | WL_cwt_sym2_32_medianabsC | wavelet,cwt | 0.000201 | 0.00617

---Outlier properties---
1689 | DN_RemovePoints_absclose_05_ac2diff | correlation,outliers | 6.98e-07 | 0.00135
1688 | DN_RemovePoints_absclose_05_ac2rat | correlation,outliers | 1.89e-06 | 0.00183
1736 | DN_RemovePoints_min_01_sumabsacfdiff | correlation,outliers | 2.71e-05 | 0.00247
1766 | DN_RemovePoints_max_01_sumabsacfdiff | correlation,outliers | 2.71e-05 | 0.00247
1746 | DN_RemovePoints_min_05_sumabsacfdiff | correlation,outliers | 5.06e-05 | 0.00308
1714 | DN_RemovePoints_absfar_01_sumabsacfdiff | correlation,outliers | 0.000259 | 0.0071

---Simulated walker---
2620 | PH_Walker_biasprop_01_05_w_std | trend | 9.67e-06 | 0.00202
2607 | PH_Walker_biasprop_05_01_sw_meanabsdiff | trend | 1.39e-05 | 0.00221
2621 | PH_Walker_biasprop_01_05_w_ac1 | trend | 1.94e-05 | 0.00221
2622 | PH_Walker_biasprop_01_05_w_ac2 | trend | 2.71e-05 | 0.00247
2633 | PH_Walker_biasprop_01_05_sw_ansarib_pval | trend | 2.71e-05 | 0.00247
2542 | PH_Walker_prop_05_w_ac2 | trend | 3.72e-05 | 0.00271
2602 | PH_Walker_biasprop_05_01_w_ac2 | trend | 3.72e-05 | 0.00271
2642 | PH_Walker_momentum_2_w_ac2 | trend | 3.72e-05 | 0.00271
2520 | PH_Walker_prop_01_w_std | trend | 5.06e-05 | 0.00308
2601 | PH_Walker_biasprop_05_01_w_ac1 | trend | 5.06e-05 | 0.00308
2541 | PH_Walker_prop_05_w_ac1 | trend | 6.78e-05 | 0.00369
2521 | PH_Walker_prop_01_w_ac1 | trend | 9.02e-05 | 0.00394
2533 | PH_Walker_prop_01_sw_ansarib_pval | trend | 9.02e-05 | 0.00394
2634 | PH_Walker_biasprop_01_05_sw_distdiff | trend | 9.02e-05 | 0.00394
2606 | PH_Walker_biasprop_05_01_w_propzcross | trend | 9.64e-05 | 0.00413
2540 | PH_Walker_prop_05_w_std | trend | 0.000119 | 0.00455
2522 | PH_Walker_prop_01_w_ac2 | trend | 0.000155 | 0.00541
2641 | PH_Walker_momentum_2_w_ac1 | trend | 0.000155 | 0.00541
2526 | PH_Walker_prop_01_w_propzcross | trend | 0.000201 | 0.00617
2562 | PH_Walker_prop_09_w_ac2 | trend | 0.000201 | 0.00617
2682 | PH_Walker_runningvar_15_50_w_ac2 | trend | 0.000201 | 0.00617
2582 | PH_Walker_prop_11_w_ac2 | trend | 0.00033 | 0.00819
2661 | PH_Walker_momentum_5_w_ac1 | trend | 0.00033 | 0.00819
2662 | PH_Walker_momentum_5_w_ac2 | trend | 0.000419 | 0.00934
1608 | PH_ForcePotential_dblwell_1_05_02_pcross | dynamicalSystem | 0.000188 | 0.00617

---Stationarity---

855 | SY_LocalGlobal_AC1_unicg500 | stationarity | 0.000155 | 0.00541

3415 | ST_LocalExtrema_n100_diffmaxabsmin | distribution,stationarity | 0.000119 | 0.00455

6201 | PP_Compare_medianf4_swss5_1 | preprocessing,raw,stationarity | 0.000201 | 0.00617

6263 | PP_Compare_rav2_swss5_1 | preprocessing,raw,stationarity | 0.000201 | 0.00617

6294 | PP_Compare_rav3_swss5_1 | preprocessing,raw,stationarity | 0.000259 | 0.0071

6325 | PP_Compare_rav4_swss5_1 | preprocessing,raw,stationarity | 0.000259 | 0.0071

1809 | CO_TranslateShape_circle_25_pts_statav4_m | correlation | 0.000259 | 0.0071

2169 | SY_DriftingMean100_max | stationarity | 0.00033 | 0.00819

1791 | CO_TranslateShape_circle_15_pts_statav3_m | correlation | 0.000419 | 0.00934

1805 | CO_TranslateShape_circle_25_pts_statav2_m | correlation | 0.000419 | 0.00934

---Nonlinear time-series analysis---

5724 | NL_TSTL_LargestLyap_n1_01_001_3_1_4_to09max | nonlinear,tstool | 0.000299 | 0.00805

5731 | NL_TSTL_LargestLyap_n1_01_001_3_1_4_vse_intercept | nonlinear,tstool | 0.000419 | 0.00934

5739 | NL_TSTL_LargestLyap_n1_01_001_3_1_4_expfit_b | nonlinear,tstool | 0.000419 | 0.00934

4777 | NL_crptool_fnn_10_2_ac_fnn7 | nonlinear,dimension,crptool | 0.00033 | 0.00819

---Other---

7694 | SY_VarRatioTest_24682468_00001111_minstat | vratiotest | 5.06e-05 | 0.00308

1807 | CO_TranslateShape_circle_25_pts_statav3_m | correlation | 9.02e-05 | 0.00394

1812 | CO_TranslateShape_circle_35_pts_std | correlation | 0.000155 | 0.00541

1890 | CO_StickAngles_y_ac1_all | correlation | 0.000155 | 0.00541

1786 | CO_TranslateShape_circle_15_pts_ones | correlation | 0.000213 | 0.00648

1949 | CO_Embed2_AngleTau_50_mean_thetaac3 | correlation,embedding | 0.000259 | 0.0071

1887 | CO_StickAngles_y_ac1_n | correlation | 0.000259 | 0.0071

1813 | CO_TranslateShape_circle_35_pts_mean | correlation | 0.000275 | 0.00749

7716 | SY_KPSStest_0_10_lagmaxstat | stationarity,hypothesistest | 0.000299 | 0.00805

1884 | CO_StickAngles_y_ac1_p | correlation | 0.000419 | 0.00934

1819 | CO_TranslateShape_circle_35_pts_fours | correlation | 0.000395 | 0.00934

Here, we list features of **PVCre-hM4Di mice** from hctsa (v0.96) with corrected p < 0.01

Features have been coarsely grouped into categories, as labeled.

Each row is of the form:

featureID | featureName | featureKeywords | pValue | pValueCorr

177 | AC_nl_025 | correlation,nonlinearautocorr | 1.95e-05 | 0.025
1089 | MD_polvar_05_4 | medical,symbolic | 2.58e-05 | 0.025
1265 | CO_AddNoise_1_gaussian_ami_at_10 | correlation,AMI,entropy | 1.18e-05 | 0.025
7257 | MF_StateSpace_n4sid_1_05_1_p1_5 | model | 1.52e-05 | 0.025
7264 | MF_StateSpace_n4sid_1_05_1_ac3 | model | 2.48e-05 | 0.025
7267 | MF_StateSpace_n4sid_1_05_1_ac3n | model | 2.48e-05 | 0.025
7385 | MF_CompareAR_1_10_05_stddiff | model | 1.52e-05 | 0.025
7386 | MF_CompareAR_1_10_05_maxdiff | model | 9.05e-06 | 0.025
6247 | PP_Compare_medianf10_htdt_ksn | preprocessing,raw | 3.15e-05 | 0.0271
139 | AC_nl_12345 | correlation,nonlinearautocorr | 9.5e-05 | 0.0272
166 | AC_nl_003 | correlation,nonlinearautocorr | 6.2e-05 | 0.0272
464 | CO_CompareMinAMI_quantiles_2_80_nunique | correlation,AMI | 6.82e-05 | 0.0272
1152 | CO_AddNoise_1_quantiles_10_ami_at_10 | correlation,AMI,entropy | 6.2e-05 | 0.0272
1227 | CO_AddNoise_1_std1_10_ami_at_5 | correlation,AMI,entropy | 6.2e-05 | 0.0272
1264 | CO_AddNoise_1_gaussian_ami_at_5 | correlation,AMI,entropy | 9.5e-05 | 0.0272
1266 | CO_AddNoise_1_gaussian_ami_at_15 | correlation,AMI,entropy | 3.97e-05 | 0.0272
2620 | PH_Walker_biasprop_01_05_w_std | trend | 9.5e-05 | 0.0272
3041 | FC_LoopLocalSimple_mean_stderr_peaksize | forecasting | 4.97e-05 | 0.0272
3961 | SC_FluctAnal_2_std_50_logi_ssr | scaling | 6.2e-05 | 0.0272
4483 | SP_Summaries_welch_rect_logarea_4_1 | spectral | 6.2e-05 | 0.0272
4493 | SP_Summaries_welch_rect_logarea_5_1 | spectral | 9.5e-05 | 0.0272
4607 | SP_Summaries_fft_logarea_4_1 | spectral | 6.2e-05 | 0.0272
4617 | SP_Summaries_fft_logarea_5_1 | spectral | 9.5e-05 | 0.0272
7263 | MF_StateSpace_n4sid_1_05_1_ac2 | model | 7.69e-05 | 0.0272
7266 | MF_StateSpace_n4sid_1_05_1_ac2n | model | 7.69e-05 | 0.0272
7378 | MF_CompareAR_1_10_05_maxv | model | 9.5e-05 | 0.0272
7384 | MF_CompareAR_1_10_05_meandiff | model | 9.5e-05 | 0.0272
174 | AC_nl_024 | correlation,nonlinearautocorr | 0.000117 | 0.0277
186 | AC_nl_113 | correlation,nonlinearautocorr | 0.000143 | 0.0277

190 | AC_nl_233 | correlation,nonlinearautocorr | 0.000143 | 0.0277

463 | CO_CompareMinAMI_quantiles_2_80_std | correlation,AMI | 0.000137 | 0.0277

1151 | CO_AddNoise_1_quantiles_10_ami_at_5 | correlation,AMI,entropy | 0.000117 | 0.0277

1267 | CO_AddNoise_1_gaussian_ami_at_20 | correlation,AMI,entropy | 0.000143 | 0.0277

4442 | SP_Summaries_welch_rect_ylogareatopeak | spectral | 0.000143 | 0.0277

4566 | SP_Summaries_fft_ylogareatopeak | spectral | 0.000143 | 0.0277

7285 | MF_StateSpace_n4sid_2_05_1_c_1 | model | 0.000143 | 0.0277

7340 | MF_StateSpace_n4sid_3_05_1_p1_5 | model | 0.000143 | 0.0277

7382 | MF_CompareAR_1_10_05_firstonmin | model | 0.000117 | 0.0277

7393 | MF_CompareAR_1_10_05_bestaic | model | 0.000143 | 0.0277

7437 | MF_armax_1_1_05_1_meanabs | model | 0.000117 | 0.0277

171 | AC_nl_013 | correlation,nonlinearautocorr | 0.000174 | 0.0314

7253 | MF_StateSpace_n4sid_1_05_1_meanabs | model | 0.000174 | 0.0314

7438 | MF_armax_1_1_05_1_stde | model | 0.000174 | 0.0314

189 | AC_nl_223 | correlation,nonlinearautocorr | 0.000309 | 0.0351

459 | CO_CompareMinAMI_quantiles_2_80_max | correlation,AMI | 0.000248 | 0.0351

466 | CO_CompareMinAMI_quantiles_2_80_modef | correlation,AMI | 0.000209 | 0.0351

488 | CO_CompareMinAMI_std2_2_80_modef | correlation,AMI | 0.000282 | 0.0351

1198 | CO_AddNoise_1_even_10_fitlinb | correlation,AMI,entropy | 0.000256 | 0.0351

1228 | CO_AddNoise_1_std1_10_ami_at_10 | correlation,AMI,entropy | 0.000309 | 0.0351

1236 | CO_AddNoise_1_std1_10_fitlinb | correlation,AMI,entropy | 0.000309 | 0.0351

1273 | CO_AddNoise_1_gaussian_fitlinb | correlation,AMI,entropy | 0.000256 | 0.0351

2520 | PH_Walker_prop_01_w_std | trend | 0.000309 | 0.0351

2602 | PH_Walker_biasprop_05_01_w_ac2 | trend | 0.000309 | 0.0351

2621 | PH_Walker_biasprop_01_05_w_ac1 | trend | 0.000309 | 0.0351

4434 | SP_Summaries_welch_rect_fpolysat_a | spectral | 0.000309 | 0.0351

4437 | SP_Summaries_welch_rect_fpolysat_rmse | spectral | 0.000309 | 0.0351

4475 | SP_Summaries_welch_rect_logarea_3_1 | spectral | 0.000256 | 0.0351

4558 | SP_Summaries_fft_fpolysat_a | spectral | 0.000309 | 0.0351

4561 | SP_Summaries_fft_fpolysat_rmse | spectral | 0.000309 | 0.0351

4599 | SP_Summaries_fft_logarea_3_1 | spectral | 0.000256 | 0.0351

6944 | NL_embed_PCA_mi_10_std | embedding,pca | 0.000309 | 0.0351

7083 | MF_steps_ahead_arma_3_1_6_rmserr_4 | model,prediction | 0.000212 | 0.0351

7084 | MF_steps_ahead_arma_3_1_6_mabserr_4 | model,prediction | 0.000309 | 0.0351

7254 | MF_StateSpace_n4sid_1_05_1_stde | model | 0.000256 | 0.0351

7255 | MF_StateSpace_n4sid_1_05_1_mms | model | 0.000212 | 0.0351

7439 | MF_armax_1_1_05_1_mms | model | 0.000309 | 0.0351

7448 | MF_armax_1_1_05_1_ac3 | model | 0.000309 | 0.0351

7451 | MF_armax_1_1_05_1_ac3n | model | 0.000309 | 0.0351

182 | AC_nl_036 | correlation,nonlinearautocorr | 0.00037 | 0.0358

462 | CO_CompareMinAMI_quantiles_2_80_mean | correlation,AMI | 0.000345 | 0.0358

490 | CO_CompareMinAMI_std2_2_80_nlocmax | correlation,AMI | 0.000346 | 0.0358

1456 | SB_TransitionpAlphabet_40_1_trfexp_b | symbolic,transitionmat | 0.00037 | 0.0358

2046 | NW_VisibilityGraph_norm_dgaussk_resAC2 | network,visibilityGraph | 0.00037 | 0.0358

2830 | EN_Randomize_statdist_ac3fexpa | entropy,slow | 0.00037 | 0.0358

2887 | EN_Randomize_dyndist_ac2fexpa | entropy,slow | 0.00037 | 0.0358

2893 | EN_Randomize_dyndist_ac3fexpa | entropy,slow | 0.00037 | 0.0358

3529 | SB_MotifTwo_diff_uuud | symbolic,motifs | 0.000342 | 0.0358

5114 | NL_TSTL_acp_mi_1__10_ac1_acpf_6 | nonlinear,correlation | 0.00037 | 0.0358

7043 | MF_steps_ahead_ar_best_6_rmserr_6 | model,prediction,arfit | 0.00037 | 0.0358

7093 | MF_steps_ahead_arma_3_1_6_meandiffrms | model,prediction | 0.00037 | 0.0358

3038 | FC_LoopLocalSimple_mean_stderr_chn | forecasting | 0.000443 | 0.0361

3522 | SB_MotifTwo_diff_duuu | symbolic,motifs | 0.000421 | 0.0361

3885 | SC_FluctAnal_2_nothing_50_logi_linfitint | scaling | 0.000443 | 0.0361

4057 | SC_FluctAnal_2_dfa_50_0_logi_ssr | scaling | 0.000443 | 0.0361

4421 | SP_Summaries_welch_rect_centroid | spectral | 0.000387 | 0.0361

4433 | SP_Summaries_welch_rect_fpoly2_rmse | spectral | 0.000443 | 0.0361

4436 | SP_Summaries_welch_rect_fpolysat_r2 | spectral | 0.000443 | 0.0361

4545 | SP_Summaries_fft_centroid | spectral | 0.000387 | 0.0361

4557 | SP_Summaries_fft_fpoly2_rmse | spectral | 0.000443 | 0.0361

4560 | SP_Summaries_fft_fpolysat_r2 | spectral | 0.000443 | 0.0361

7346 | MF_StateSpace_n4sid_3_05_1_ac2 | model | 0.000443 | 0.0361

7347 | MF_StateSpace_n4sid_3_05_1_ac3 | model | 0.000443 | 0.0361

7350 | MF_StateSpace_n4sid_3_05_1_ac3n | model | 0.000443 | 0.0361

7441 | MF_armax_1_1_05_1_p1_5 | model | 0.000443 | 0.0361

7768 | CP_ML_StepDetect_l1pwc_10_nsegments | stepdetection | 0.000414 | 0.0361

3517 | SB_MotifTwo_diff_ddud | symbolic,motifs | 0.000459 | 0.0366

3688 | SB_MotifThree_quantile_caab | symbolic,motifs | 0.000458 | 0.0366

750 | SC_fastdfa_exponent | dfa,scaling,mex | 0.000528 | 0.0379

1190 | CO_AddNoise_1_even_10_ami_at_10 | correlation,AMI,entropy | 0.000528 | 0.0379

1197 | CO_AddNoise_1_even_10_fitlina | correlation,AMI,entropy | 0.000528 | 0.0379

1411 | SB_TransitionMatrix_3ac_maxeigcov | symbolic,transitionmat | 0.000528 | 0.0379

1945 | CO_Embed2_AngleTau_50_min_thetaac1 | correlation,embedding | 0.000528 | 0.0379

3641 | SB_MotifThree_quantile_aacc | symbolic,motifs | 0.0005 | 0.0379

4151 | SC_FluctAnal_2_dfa_50_1_2_logi_se1 | scaling | 0.000528 | 0.0379

4152 | SC_FluctAnal_2_dfa_50_1_2_logi_se2 | scaling | 0.000528 | 0.0379

7025 | MF_steps_ahead_ar_2_6_maxdiffrms | model,prediction | 0.000528 | 0.0379

7047 | MF_steps_ahead_ar_best_6_meandiffrms | model,prediction,arfit | 0.000528 | 0.0379

7048 | MF_steps_ahead_ar_best_6_maxdiffrms | model,prediction,arfit | 0.000528 | 0.0379

1235 | CO_AddNoise_1_std1_10_fitlina | correlation,AMI,entropy | 0.000628 | 0.0389

1406 | SB_TransitionMatrix_3ac_sumdiagcov | symbolic,transitionmat | 0.000628 | 0.0389

1887 | CO_StickAngles_y_ac1_n | correlation | 0.000628 | 0.0389

2282 | FC_Surprise_T1_50_3_q_500_mean | information,symbolic | 0.000628 | 0.0389

2642 | PH_Walker_momentum_2_w_ac2 | trend | 0.000628 | 0.0389

3381 | ST_LocalExtrema_n50_meanmax | distribution,stationarity | 0.000628 | 0.0389

4006 | SC_FluctAnal_2_rsrange_50_logi_alpha | scaling | 0.000628 | 0.0389

6241 | PP_Compare_medianf10_kscn_adiff | preprocessing,raw | 0.000628 | 0.0389

6939 | NL_embed_PCA_mi_10_perc_1 | embedding,pca | 0.000628 | 0.0389

6945 | NL_embed_PCA_mi_10_range | embedding,pca | 0.000628 | 0.0389

6947 | NL_embed_PCA_mi_10_top2 | embedding,pca | 0.000628 | 0.0389

7011 | MF_steps_ahead_ar_2_6_rmserr_3 | model,prediction | 0.000628 | 0.0389

7040 | MF_steps_ahead_ar_best_6_rmserr_5 | model,prediction,arfit | 0.000628 | 0.0389

7086 | MF_steps_ahead_arma_3_1_6_rmserr_5 | model,prediction | 0.000628 | 0.0389

7463 | MF_armax_2_2_05_1_AR_2 | model | 0.000628 | 0.0389

7490 | MF_armax_2_2_05_1_ftbth | model | 0.000568 | 0.0389

7706 | SY_KPSStest_0_stat | stationarity,hypothesistest | 0.000628 | 0.0389

460 | CO_CompareMinAMI_quantiles_2_80_range | correlation,AMI | 0.000673 | 0.0413

1051 | DK_crinkle_statistic | misc | 0.000744 | 0.0414

1413 | SB_TransitionMatrix_3ac_stdeigcov | symbolic,transitionmat | 0.000744 | 0.0414

2769 | EN_mse_1-10_2_015_diff1_sampen_s6 | entropy,sampen,controlen,mse | 0.000744 | 0.0414

3613 | SB_MotifThree_quantile_acc | symbolic,motifs | 0.000705 | 0.0414

4240 | SC_FluctAnal_sign_2_dfa_50_2_logi_r2_se1 | scaling | 0.000744 | 0.0414

4430 | SP_Summaries_welch_rect_fpoly2csS_p3 | spectral | 0.000744 | 0.0414

4435 | SP_Summaries_welch_rect_fpolysat_b | spectral | 0.000744 | 0.0414

4554 | SP_Summaries_fft_fpoly2csS_p3 | spectral | 0.000744 | 0.0414

4559 | SP_Summaries_fft_fpolysat_b | spectral | 0.000744 | 0.0414

6955 | NL_embed_PCA_mi_10_fb01 | embedding,pca | 0.000721 | 0.0414

7088 | MF_steps_ahead_arma_3_1_6_ac1_5 | model,prediction | 0.000744 | 0.0414

7408 | MF_armax_3_1_05_1_p1_5 | model | 0.000744 | 0.0414

7708 | SY_KPSStest_1_stat | stationarity,hypothesistest | 0.000744 | 0.0414

242 | CO_HistogramAMI_even_5_1 | information,correlation,AMI | 0.000878 | 0.0459

433 | IN_AutoMutualInfoStats_diff_20_kraskov1_4_stdami | information,correlation,AMI | 0.000878 | 0.0459
1160 | CO_AddNoise_1_quantiles_10_fitlinb | correlation,AMI,entropy | 0.000878 | 0.0459
1189 | CO_AddNoise_1_even_10_ami_at_5 | correlation,AMI,entropy | 0.000878 | 0.0459
1193 | CO_AddNoise_1_even_10_fitexpa | correlation,AMI,entropy | 0.000878 | 0.0459
2542 | PH_Walker_prop_05_w_ac2 | trend | 0.000878 | 0.0459
2601 | PH_Walker_biasprop_05_01_w_ac1 | trend | 0.000878 | 0.0459
7026 | MF_steps_ahead_ar_2_6_stddiffrms | model,prediction | 0.000878 | 0.0459
7044 | MF_steps_ahead_ar_best_6_mabserr_6 | model,prediction,arfit | 0.000878 | 0.0459
4418 | SP_Summaries_welch_rect_wmax_5 | spectral | 0.000911 | 0.047
4542 | SP_Summaries_fft_wmax_5 | spectral | 0.000911 | 0.047
160 | AC_nl_033 | correlation,nonlinearautocorr | 0.00103 | 0.0471
183 | AC_nl_046 | correlation,nonlinearautocorr | 0.00103 | 0.0471
247 | CO_HistogramAMI_even_10_1 | information,correlation,AMI | 0.00103 | 0.0471
1272 | CO_AddNoise_1_gaussian_fitlina | correlation,AMI,entropy | 0.00103 | 0.0471
2521 | PH_Walker_prop_01_w_ac1 | trend | 0.00103 | 0.0471
2540 | PH_Walker_prop_05_w_std | trend | 0.00103 | 0.0471
2560 | PH_Walker_prop_09_w_std | trend | 0.00103 | 0.0471
2622 | PH_Walker_biasprop_01_05_w_ac2 | trend | 0.00103 | 0.0471
2746 | EN_SampEn_5_02_diff1_meanchsampen | entropy,sampen,controlen | 0.00103 | 0.0471
2764 | EN_mse_1-10_2_015_diff1_sampen_s1 | entropy,sampen,controlen,mse | 0.00103 | 0.0471
3636 | SB_MotifThree_quantile_aaba | symbolic,motifs | 0.00102 | 0.0471
3863 | MF_AR_arcov_5_a2 | model,fit | 0.00103 | 0.0471
3866 | MF_AR_arcov_5_a5 | model,fit | 0.00103 | 0.0471
3875 | DN_SimpleFit_sin1_resAC2 | model | 0.00103 | 0.0471
4190 | SC_FluctAnal_2_dfa_50_1_3_logi_r2_linfitint | scaling | 0.00103 | 0.0471
4420 | SP_Summaries_welch_rect_wmax_25 | spectral | 0.000991 | 0.0471
4544 | SP_Summaries_fft_wmax_25 | spectral | 0.000991 | 0.0471
6245 | PP_Compare_medianf10_kscn_relent | preprocessing,raw | 0.00103 | 0.0471
7012 | MF_steps_ahead_ar_2_6_mabserr_3 | model,prediction | 0.00103 | 0.0471
7080 | MF_steps_ahead_arma_3_1_6_rmserr_3 | model,prediction | 0.00103 | 0.0471
3642 | SB_MotifThree_quantile_abaa | symbolic,motifs | 0.00105 | 0.0477
7774 | CP_ML_StepDetect_l1pwc_10_meanstepint | stepdetection | 0.00107 | 0.0481
145 | AC_nl_135 | correlation,nonlinearautocorr | 0.00142 | 0.0489
224 | CO_HistogramAMI_std2_2_3 | information,correlation,AMI | 0.00121 | 0.0489
264 | CO_HistogramAMI_quantiles_10_3 | information,correlation,AMI | 0.00121 | 0.0489

312 | IN_AutoMutualInfoStats_40_gaussian_modeperiodmax | information,correlation,AMI | 0.00131 | 0.0489
455 | CO_CompareMinAMI_std1_2_80_modef | correlation,AMI | 0.00126 | 0.0489
468 | CO_CompareMinAMI_quantiles_2_80_nlocmax | correlation,AMI | 0.00115 | 0.0489
981 | HT_DistributionTest_chi2beta50 | hypothesistest,distribution,raw | 0.00121 | 0.0489
1108 | CO_tc3_3_denom | correlation,nonlinear | 0.00142 | 0.0489
1231 | CO_AddNoise_1_std1_10_fitexpa | correlation,AMI,entropy | 0.00142 | 0.0489
1395 | SB_TransitionMatrix_3ac_T3 | symbolic,transitionmat | 0.00137 | 0.0489
1425 | SB_TransitionMatrix_4ac_mineig | symbolic,transitionmat | 0.00142 | 0.0489
1714 | DN_RemovePoints_absfar_01_sumabsacfdiff | correlation,outliers | 0.00142 | 0.0489
2522 | PH_Walker_prop_01_w_ac2 | trend | 0.00142 | 0.0489
2541 | PH_Walker_prop_05_w_ac1 | trend | 0.00121 | 0.0489
2624 | PH_Walker_biasprop_01_05_w_min | trend | 0.00142 | 0.0489
2641 | PH_Walker_momentum_2_w_ac1 | trend | 0.00121 | 0.0489
2778 | EN_mse_1-10_2_015_diff1_meanSampEn | entropy,sampen,controlen,mse | 0.00142 | 0.0489
3010 | FC_LocalSimple_mean3_stderr | forecasting | 0.00142 | 0.0489
3607 | SB_MotifThree_quantile_aac | symbolic,motifs | 0.00116 | 0.0489
3808 | SB_MotifThree_diffquant_caab | symbolic,motifs | 0.00134 | 0.0489
3856 | MF_AR_arcov_4_a4 | model,fit | 0.00142 | 0.0489
3865 | MF_AR_arcov_5_a4 | model,fit | 0.00142 | 0.0489
3874 | DN_SimpleFit_sin1_resAC1 | model | 0.00121 | 0.0489
3974 | SC_FluctAnal_2_std_50_logi_r2_linfitint | scaling | 0.00121 | 0.0489
4070 | SC_FluctAnal_2_dfa_50_0_logi_r2_linfitint | scaling | 0.00142 | 0.0489
4110 | SC_FluctAnal_2_dfa_50_2_logi_meanssr | scaling | 0.00142 | 0.0489
4166 | SC_FluctAnal_2_dfa_50_1_2_logi_r2_linfitint | scaling | 0.00142 | 0.0489
4223 | SC_FluctAnal_sign_2_dfa_50_2_logi_se1 | scaling | 0.00142 | 0.0489
4224 | SC_FluctAnal_sign_2_dfa_50_2_logi_se2 | scaling | 0.00142 | 0.0489
4417 | SP_Summaries_welch_rect_tau | spectral | 0.00126 | 0.0489
4431 | SP_Summaries_welch_rect_fpoly2_sse | spectral | 0.00121 | 0.0489
4541 | SP_Summaries_fft_tau | spectral | 0.00126 | 0.0489
4555 | SP_Summaries_fft_fpoly2_sse | spectral | 0.00121 | 0.0489
5726 | NL_TSTL_LargestLyap_n1_01_001_3_1_4_to07max | nonlinear,tstool | 0.00141 | 0.0489
6463 | MF_GP_hyperparameters_covSEiso_covNoise_1_200_resample_rmserr | model,GaussianProcess | 0.00142 | 0.0489
6481 | MF_GP_hyperparameters_covSEiso_covPeriodic_covNoise_1_200_resample_minS | model,GaussianProcess | 0.00142 | 0.0489
6743 | WL_dwtcoeff_sym2_5_maxd_l2 | wavelet,dwt | 0.00142 | 0.0489

6940 | NL_embed_PCA_mi_10_perc_2 | embedding,pca | 0.00142 | 0.0489

7037 | MF_steps_ahead_ar_best_6_rmserr_4 | model,prediction,arfit | 0.00142 | 0.0489

7041 | MF_steps_ahead_ar_best_6_mabserr_5 | model,prediction,arfit | 0.00121 | 0.0489

7089 | MF_steps_ahead_arma_3_1_6_rmserr_6 | model,prediction | 0.00121 | 0.0489

7091 | MF_steps_ahead_arma_3_1_6_ac1_6 | model,prediction | 0.00142 | 0.0489

7095 | MF_steps_ahead_arma_3_1_6_stddiffrms | model,prediction | 0.00142 | 0.0489

7258 | MF_StateSpace_n4sid_1_05_1_p2_5 | model | 0.00121 | 0.0489

7262 | MF_StateSpace_n4sid_1_05_1_ac1 | model | 0.00142 | 0.0489

7265 | MF_StateSpace_n4sid_1_05_1_ac1n | model | 0.00142 | 0.0489

7269 | MF_StateSpace_n4sid_1_05_1_acsnd0 | model | 0.00142 | 0.0489

7272 | MF_StateSpace_n4sid_1_05_1_dwts | model | 0.00142 | 0.0489

7275 | MF_StateSpace_n4sid_1_05_1_sbc1 | model | 0.00121 | 0.0489

7370 | MF_CompareAR_1_10_all_maxdiff | model | 0.00142 | 0.0489

7466 | MF_armax_2_2_05_1_maxda | model | 0.00142 | 0.0489

7701 | SY_PPtest_0_5_ar_t1_meanstat | unitroot | 0.00142 | 0.0489

7779 | CP_ML_StepDetect_l1pwc_10_medianstepint | stepdetection | 0.00112 | 0.0489
